# Monandrous flies do remate: plastic and evolutionary consequences of heat exposure on mating behaviour and fertility in *Drosophila subobscura*

**DOI:** 10.1101/2024.11.12.623111

**Authors:** Afonso Grandela, Marta A. Antunes, Marta A. Santos, Ana S. Quina, Isabel R. Costa, Guilherme Borges, Margarida Matos, Leonor R. Rodrigues, Pedro Simões

## Abstract

Recent work has reinforced the importance of fertility in predicting population persistence under global warming, especially male fertility. However, most studies on mating and reproductive performance have used as model organism polyandrous species, remaining unclear if and how will monandrous species respond. This is unfortunate as one could expect monandrous species to be particularly susceptible, given their inability to use sperm from multiple males to prevent reduced progeny production due to male sterility. It could be that monandrous females can overcome this toll in fertility by plastically changing their mating rate, or that monandrous populations evolve the ability to remate upon repeated exposure to heat. To test this, we studied the real-time evolution as well as the thermal plastic response of two geographically distinct populations of the monandrous species *Drosophila subobscura* after 45 generations evolving under a warming scenario. We assessed the impact of heat on male mating behaviour and fertility, as well as the impact of reduced male fertility on female mating behaviour and fertility. We found that in males the toll in mating behaviour due to heat can be recovered with time, whereas that in reproductive success cannot. On the other hand, females exposed to heat-stressed males significantly increased their remating rate, allowing to recover considerably their own reproductive success. Interestingly, most responses were plastic, with no striking effect of adaptation to warming nor of geographical origin. Our work brings new insights into the effects of heat on monandrous species and establishes a model to study how a shift in mating system may affect species’ ability to respond to climate change.

## Introduction

Rising temperatures have intensified in the last 100 years, with recent climatic models pointing to an even higher increase, ranging between 1.4°C and 4.2°C in average global temperature, as well as an intensification in frequency and duration of heatwave events until the end of this century (IPCC 2022). These environmental shifts have detrimental effects on biodiversity worldwide, influencing species’ habitat distributions and leading to population extinction (Nunez et al., 2019; Pecl et al., 2017).

In the past, studies have mostly focused on the effects of rising temperatures on the upper critical thermal limit of individuals. However, recently, it has been reinforced that different traits can have contrasting thermal sensitivities, with fertility being negatively affected at lower temperatures than other traits, such as viability and survival (Ma et al., 2021; Parratt et al., 2021; Walsh et al., 2019; Zhang et al., 2013). Thermal sensitivity has also been shown to differ between sexes, with males often being more affected by rising temperatures than females (David et al., 2005; Iossa, 2019; Sales et al., 2018; Zwoinska et al., 2020). Accordingly, males’ thermal fertility limits seem to be the best predictors of species’ persistence and distribution in a global warming scenario (Parratt et al., 2021; van Heerwaarden and Sgrò, 2021).

In exploring the impact of rising temperatures on fertility, one important aspect to consider is whether heat-induced sterility in males is transient or permanent during the individuals’ lifetime. Studies tackling this question show contrasting results, some reporting a partial to total recovery of male fertility after heat stress in both the developmental (Canal Domenech and Fricke, 2023, 2022; Sales et al., 2021; Walsh et al., 2021) and the adult stages (Canal Domenech and Fricke, 2023; Parrett et al., 2024; Sales et al., 2021), while others find permanent or long-lasting sterility following heat exposure during adulthood (Meena et al., 2024; Parratt et al., 2021; Walsh et al., 2021). In addition, both pre and postcopulatory mating behaviours can be simultaneously affected by rising temperatures (Costa et al., 2023; Farrow et al., 2022; Leith et al., 2020; Pilakouta et al., 2023; Sutter et al., 2019; Vasudeva et al., 2021), and are important determinants of organisms’ ability to cope with thermal stress (Gómez-Llano et al., 2021; Sutter et al., 2019; Vasudeva et al., 2021). For instance, mating rate has been shown to change with heat, being advantageous for females to mate with multiple males when male heat-induced sterility is prevalent; this is because mating multiply can improve female reproductive success (Sutter et al., 2019; Vasudeva et al., 2021), functioning as a buffer against population decline. Unfortunately, the great majority of studies have been done in polyandrous species and neglect the possible impact of the mating system on the ability of organisms to respond to rising temperatures (but see Baur et al., 2024; Moiron et al., 2022).

The mating system is at the very basis of describing a species reproductively and it is well established how it can shape interactions between sexes (e.g., the intensity of sexual conflict; Holman and Kokko, 2013, within sexes (e.g., male-male competition; Lizé et al., 2012a), as well as populations’ ecology and evolutionary potential (e.g., inbreeding and effective population size (Holman and Kokko, 2013). In the context of global warming, one could expect strictly monandrous species to be particularly susceptible. For instance, by definition, monandrous females, unlike polyandrous females, will only mate once and, as such, will necessarily suffer the drop in offspring production imposed by male heat-induced sterility. This toll on female fertility could be overcome if the female mating rate is plastic, in general, or at least if monandrous populations exposed to heat evolve the ability to remate. Yet, given the general lack of studies in monandrous species, it is hard to predict if and which of these outcomes is the most likely.

Interestingly, while plastic responses to heat are frequently studied, knowledge on the genetic adaptation of organisms to global warming is lagging behind (Kellermann and van Heerwaarden, 2019). Out of those that do look at the adaptive potential, most demonstrate that species fail to show evolutionary responses to warming environments (Kellermann et al., 2009; Kinzner et al., 2019; Schou et al., 2014; van Heerwaarden and Sgrò, 2021). Still, recent work performing experimental evolution in Drosophila has found adaptive responses to warming (Santos et al., 2023), showing that distinct populations can respond differently (Antunes et al., 2024; Santos et al., 2023). Furthermore, indirect evidence from studies comparing the response of different populations supports the idea that intraspecific variation plays a role in shaping thermal responses (Austin and Moehring, 2019; Porcelli et al., 2017). It is undeniable that more studies of adaptation to warming are needed to accurately predict population persistence. In particular, understanding the adaptive potential of monandrous species to warming will elucidate on how mating interactions affect populations’ evolutionary response to thermal conditions (Edelsparre et al., 2024), a topic about which we know very little.

Taking all of this into consideration, understanding if and how monandrous species respond to heat stress will be key to accurately predict and eventually mitigate the impacts of global warming on biodiversity. To fill this gap, we used the commonly monandrous fruit fly *Drosophila subobscura* (Fisher et al., 2013). *D. subobscura* is an exceptional species to study thermal adaptation, due to its wide geographical distribution and chromosomal inversions that seem to respond to global warming (Balanyá et al., 2006) and even heatwave occurrences (Rodríguez-Trelles et al., 2013). In fact, it is known that temperature changes lead to both plastic (Santos et al., 2021a; Simões et al., 2020) as well as evolutionary responses in this species (Mesas et al., 2021; Santos et al., 2023). Furthermore, there is evidence of multiple mating (ca. 14%) under certain circumstances, indicating that some genetic variation for mating behaviour exists within this species (Fisher et al., 2013).

Here, we used two sets of laboratorial populations of *D. subobscura* founded from different latitudes, one from low-latitude (Portugal) and other from high-latitude (The Netherlands; Simões et al., 2017), subjected to a warming scenario for 45 generations after lab adaptation. We previously observed evolution of increased reproductive output in the low-latitude populations after 39 generations under these warming conditions (Santos et al., 2023). We also observed clear differences in the genetic background of these populations, with diverse transcriptomic responses after 23 generations of thermal evolution (Antunes et al., 2024). By exposing evolved males to a sub-lethal heatwave treatment during the early stages of adulthood, our goals were to know: 1) how are male mating behaviour and fertility influenced by heat and whether this effect is transient or not; 2) what are the impacts of heat-induced male sterility on female mating behaviour and fertility; and 3) if thermal selection and population geographical origin play a role in how populations respond.

## Material and methods

### Experimental populations & Selective regimes

This study involved two populations of *D. subobscura*, one from Portugal and another from The Netherlands (previously labelled as “PT” and “NL”, respectively, see Simões et al., 2017). These populations were three-fold replicated by generation four in the lab and were kept under well-defined conditions for 70 generations before the onset of the thermal experimental evolution, including: discrete 28-day generations, at a constant temperature of 18°C under a 12:12 light:dark cycle; controlled density in the adult (40 individuals per vial) and developmental stage (70 eggs per vial), with egg-collection to the next generation from 6-10 day old imago flies, in large population sizes (500-1000 individuals; Simões et al., 2017). To study thermal adaptation to warming, each of these populations was divided and placed under three different selection regimes, differing in their temperature cycle: the Control regime, maintained at a constant temperature of 18°C; the Fluctuating regime, maintained with a fixed circadian temperature daily cycle ranging between 15°C and 21°C and a mean daily temperature also of 18°C; and the Warming regime, maintained with a progressive increase in thermal average and amplitude (an increase of 0.18°C and 0.54°C each generation, respectively), starting with a circadian temperature daily cycle as the Fluctuating regime (see Santos et al., 2021b and Figure S1). Populations from the Warming regime severely crashed when the maximum temperature reached 30.2 °C (at generation 22). For that reason, in the 24^th^ generation, the thermal cycle in this regime was reversed to that of the 20^th^ generation and maintained from then onwards, oscillating between 13.5°C and 29.4°C with a mean temperature of 21.4°C (Santos et al., 2023). Populations were kept under this constant cycle for 21 more generations, i.e. they were at their 45^th^ generation since the start of the thermal regimes when the assays described below were performed. Here, we focused on populations from the Control and Warming regimes, to understand the populations’ responses to global warming, while the males and females from the Fluctuating regime were used as a second source of control in some assays (see details below).

### Experimental assays

#### Recovery of male mating behaviour and fertility following heatwave

To assess the long-term consequences of high temperatures on male mating behaviour and fertility we monitored male performance from the 4^th^ to the 12^th^ day of adult life after exposure to a heatwave (see protocol below). To remove parental and grand-parental environmental effects (‘common garden’ protocol), 18 vials (with approximately 70 eggs each) from the Control and Warming populations were maintained in the environment of the control regime (hereafter, “the control environment”) for two generations.

Virgin males recently emerged as adults from the 3^rd^ generation in the control environment were anaesthetised with CO_2_ and maintained at 18°C in groups of five. A subset of the isolated males was maintained in these conditions (“non-stressed” treatment) until the beginning of the assay. The remaining virgin males, one day after emergence, were exposed to 31°C for 69 hours, followed by a 4h recovery period at 18°C (“stressed males” treatment). Exposure to such a peak temperature is expected to be quite frequent by the end of the century in Europe, from where both populations originate, with the mean temperature being predicted to reach values above 30°C for 40 to 60 days per year (Carvalho et al., 2021). The imposed heat conditions inflicted a highly negative effect on male fertility (ca. 59.4% drop in fertility after one mating compared to non-stressed males; see Figure 2), without significantly compromising mortality (i.e., below 30%).

After the recovery period, both stressed and non-stressed males were paired with virgin females from the Fluctuating regime for approximately 28 hours. These females were directly taken from the Fluctuating regime (without a ‘common garden’ protocol) immediately after emergence and maintained in groups of ten at 18°C for 4 to 6 days before being paired. By using non-stressed females from the Fluctuating populations, differences in responses between treatments can be assigned exclusively to differences in males, allowing the exclusion of a female effect or of those effects stemming from males and females co-evolving within selection regime that would be masked otherwise.

In the first two hours of each mating assay, mating behaviour was continuously observed, with the beginning of courtship, and the beginning and end of the first copulation being registered. This allowed to estimate courtship latency (time elapsed between pairing and beginning of male courtship), copulation latency (time elapsed between pairing and beginning of copulation), and copulation duration (time elapsed between the beginning and the end of the copula). In addition, in the following 9 hours, the occurrence of matings was checked every 5 minutes. The observations were done at room temperature (21°C), after which the couples were placed in an incubator at 18°C. At the end of the mating period, males were moved to a new vial kept at 18°C, while females were kept in the same vial where they laid eggs for 44 hours. To assess reproductive success (number of offspring produced) the number of emerged adults was counted seven days after the emergence of the first adult. Other fertility traits, such as fecundity (number of eggs) and offspring viability (% of eggs giving rise to adults) were also estimated (see Supplement 1). Three and eight days after the first mating, the same males had the possibility to mate a second and a third time, respectively, with new virgin females in separate vials (see Figure S2). Reproductive success and mating behaviour were assessed after these two additional matings following a protocol similar to the one described for the first mating.

A total of 24 males per treatment (Stressed and Non-stressed), per replicate population (1, 2, and 3), per history (Portuguese and Dutch), and per selection regime (Control and Warming) were tested (576 males in total). The assays used an experimental full-block design performed on 3 consecutive days, with same numbered Control and Warming replicate populations being tested on the same day (blocks 1, 2 and 3).

#### Female mating behaviour and fertility following male heatwave

To understand the impact of reduced male fertility on female mating behaviour and fertility, three distinct treatments were designed where females were paired twice with non-stressed and/or stressed males (see Figure S3): females had the opportunity to mate consecutively with 1) two non-stressed males (Non-stressed + non-stressed treatment); 2) one stressed male and one non-stressed male (Stressed + non-stressed treatment); 3) two stressed males (Stressed + stressed treatment). A protocol similar to the one described for the first assay was followed to obtain stressed males and non-stressed males and females, with the exception that Control and Warming populations were maintained in the control environment (at 18°C) for a single generation before being allocated to one of the two thermal treatments. Following the same reasoning as in the previous assay, females were crossed with virgin males from the Fluctuating regime.

Virgin females were paired for 28 hours with two 5-day-old males, the first when they were 4-day-old and the second 72 hours later. As in the previous assay, the courtship and copulation latencies and the copulation duration were obtained for the first 2 hours of each mating opportunity and mating occurrence was checked every 5 minutes, this time for approximately 10 more hours. As before, females remained for 44 hours in each vial after male removal, and the number of adult offspring produced (as well as the number of eggs and the egg to adult viability; see Supplement 1) was registered for each mating opportunity. Note that we considered that a female remated when both the first and second copulations were observed or when a female produced adult offspring after being paired with the first male and then was observed copulating with the second male.

A total of 24 females per treatment (Non-stressed + non-stressed, Stressed + non-stressed, and Stressed + stressed), per replicate population (1, 2, and 3), per history (Portuguese and Dutch), and per selection regime (Control and Warming) were tested (864 females in total). The experiment was blocked as described in the previous assay.

### Statistical analyses

All statistical analyses were performed using R (R Core Team, 2022, version 2022.7.1.554). Linear models (LM) and linear mixed-effect models (LMM) using the package *lme4* (Bates et al., 2015), mixed-effects Cox models using the package *coxme* (Terry, 2022) and generalized mixed-effects models (GLMM) with *glmmTMB* (Brooks et al., 2017) were performed depending on the dependent variables and the error structure of the data (see Tables S1 and S2). A “sum to zero” contrast option was defined for each factor. Raw individual data was used in the analysis of all dependent variables. Originally, models included interactions between fixed and random factors, but they were dropped since the inclusion of these terms led to convergency issues and or reduced the fit of the models. Maximal models were simplified by removing non-significant interaction terms from the highest- to the lowest-order interaction (see Tables S1 and S2). Model simplification ended when the lowest AIC (Akaike information criterion) value was reached. The significance level of different factors and their interactions were obtained through analyses of variance (ANOVA Type III; package *car*; Fox & Weisberg, 2019). Importantly, when the best model contained significant triple interactions or more than one double interaction, additional analyses were performed by subsetting the original dataset into the different levels of a fixed factor present in the model (Tables S1 and S2). When double interactions were present in minimal models, the *emmeans* package (Length, 2020) was used to perform *a posteriori* contrasts using Tukey tests. All graphical representations were done with the mean values of each replicate population and generated with *ggplot2* package (Wickham, 2016).

#### Recovery of male mating behaviour and fertility following heatwave

To analyse how a heatwave during early life influenced males’ mating behaviour and reproductive output, four traits were studied: Courtship latency, copulation latency and duration, and reproductive success (see Table S1).

Courtship latency and copulation latency were analysed using a mixed-effect Cox model with a gaussian error distribution, with the data considered censored whenever copulation was not observed within the time established for the mating assay. The data was box cox transformed (courtship latency: λ = 0.03; copulation latency λ = 0.75; *MASS* package; Venables & Ripley, 2002) to improve the fit of the models. A LMM model with a gaussian error distribution was used to analyse the copulation duration after the data was box cox transformed (copulation duration: λ = 1). The models for these three traits included History (Dutch or Portuguese), Selection (Control or Warming), Treatment (Stressed male or Non-stressed male), Mating Opportunity (First, Second, or Third), and all possible interactions as fixed categorical factors. Block, defined as the set of same numbered replicate populations (1, 2, or 3), was included as a random factor. Given that each male had three mating opportunities, we intended to account for repeated measures (Park et al., 2009) by adding the interaction between male ID (unique identification of each male) and Mating Opportunity as a covariate in the model. However, due to convergence issues, this covariate was excluded from the analysis.

To assess reproductive success (number of adult offspring) a GLMM model with a quasi-Poisson error distribution and a parameter to account for zero-inflated data (ziformula ∼1; package *glmmTMB*) was used. The model included the same fixed and random factors described above. In addition to those, we accounted for repeated measures by adding the interaction between male ID (unique identification of each male) and Mating Opportunity as a covariate in the model. Due to a significant quadruple interaction between all factors affecting reproductive success (see Table S3), analyses were done separately for populations from different bio-geographical origins (factor History). The analysis was similar to that described above but without History as fixed factor. A GLMM model with a quasi-Poisson error distribution and another with a Poisson error distribution were used to analyse the data from Portuguese and Dutch populations respectively, both models including a parameter to account for zero-inflated data (ziformula ∼1; package *glmmTMB*)

#### Female mating behaviour and fertility following male heatwave

To analyse how female mating behaviour and fertility were affected by male thermal treatment (stressed or non-stressed), five traits were studied: courtship latency, copulation latency and duration, female propensity to remate, and reproductive success (see Table S2).

To analyse courtship and copulation latencies mixed-effect Cox models with a gaussian error distribution were used, with the data considered censored whenever copulation was not observed within the time established for the mating assay. The data was box cox transformed (courtship latency: λ = 0.27; copulation latency λ = 0.58; *MASS* package) prior to the analysis. A LMM model with a gaussian error distribution applied to box cox transformed data (λ = 0.45) was used to analyse copulation duration. The models for these three traits used the fixed factors History (Dutch or Portuguese), Selection (Control or Warming), Treatment (Non-stressed + non-stressed, Stressed + non-stressed or Stressed + stressed), Mating Opportunity (First or Second) and their interactions. Block was added as a random factor. Following the same rationale as in the assay targeting males, given that the same females were tested in a first and second mating opportunities, we intend to account for repeated measures by adding the interaction between female ID (unique identification of each female) and Mating Opportunity as a covariate in the model. Once again, due to convergence issues, this covariate was excluded from the analysis.

Reproductive success was analysed using a GLMM model with a quasi-Poisson error distribution and a parameter to account for zero inflation (ziformula ∼1) and included the independent variables and all possible interactions described above. In addition to those, we accounted for repeated measures by adding the interaction between female ID (unique identification of each female) and Mating Opportunity as a covariate in the model. Due to multiple significant interactions involving the factor History in the analyses of reproductive success (see Table S4), new analyses were done separately for populations with distinct historical origins. The models were identical to those described above but excluding History as fixed factor.

The female propensity to remate (presence/absence of a second copulation) was analysed using a GLMM model with a binomial (Bernoulli) error distribution. This model included the same independent variables described above except for Mating Opportunity and female ID and all possible interactions.

Lastly, to confirm that remating behaviour was responsible (at least in part) for differences in reproductive success, an additional analysis of reproductive success was performed. The first model included History, Selection, Remating (No Remating or Remated), Treatment, Mating Opportunity, and their interactions as fixed factors, Block as a random factor, and the interaction between female ID and Mating Opportunity as a covariate. This variable was analysed using a GLMM model with a quasi-Poisson error distribution and a parameter to account for zero inflation (ziformula ∼1). However, convergence issues arose due to the high complexity of the model. For that reason, and because there was no effect of History nor Selection in the analysis of female propensity to remate, these factors were excluded from this analysis. Therefore, the model was similar to the one described above, excluding these two factors. Due to a significant triple interaction between Remating, Mating Opportunity and Treatment (see Table S5), further analyses were performed separating the three different treatments. The models were similar to the one including all factors but Treatment. The Non-stressed + non-stressed treatment was analysed using a GLMM model with a negative binomial error distribution, while both the Stressed + non-stressed and the Stressed + stressed treatments were analysed using models with a quasi-Poisson error distribution. All models included a parameter to account for zero-inflated data (ziformula ∼1).

## Results

### Recovery of male mating behaviour and fertility following heatwave

#### Effect of heatwave on male mating behaviour

Courtship and copulation latencies showed similar responses, both being affected by an interaction between Treatment and Mating Opportunity (*X^2^*= 37.469, p < 0.001; *X^2^* = 33.060, p < 0.001, respectively; Table S6A, Figure 1A and 1B) but not by Selection or History. Indeed, both stressed and non-stressed males showed the longest courtship and copulation latencies in their first mating, with stressed males being strikingly slower to initiate courtship and copulation than non-stressed males; however, non-stressed males took the same time before starting to court and copulate with the second and third female they were offered, while heatwave males were increasingly faster at initiating courtship and mating (see Table S7; Figure 1A and 1B). These results indicate that males exposed to heatwave were able to recover a relevant part of their behavioural performance throughout the experiment (ca. 80% and 70% shorter courtship and copulation latencies, respectively, comparing their first and third mating opportunities), although stressed males still took significantly longer to court and mate relative to non-stressed males by the third mating opportunity (Figures 1A and 1B, Table S7).

**Figure 1.**
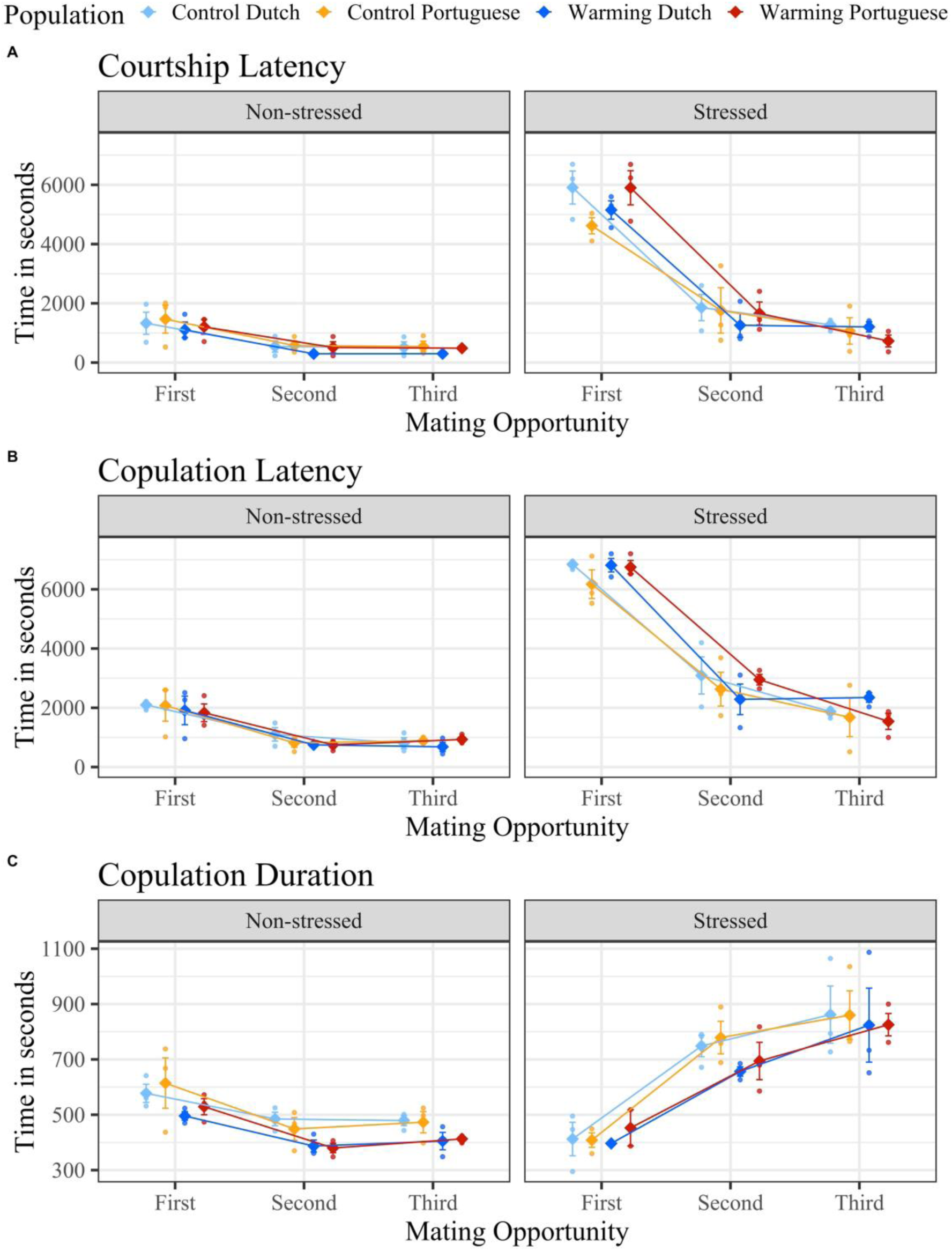
– Effect of heatwave during male adulthood on male mating behaviour. A) Courtship Latency: Time elapsed between pairing and the beginning of male courtship; B) Copulation latency: Time elapsed between pairing and the beginning of copulation; C) Copulation Duration: Time elapsed between the beginning of the copula and its ending. Colder colours represent Dutch populations, while warmer ones represent Portuguese populations. Lighter tones represent the Control regime, while darker tones represent the Warming regime. The small circles represent the mean values of each replicate population, and the big diamonds represent the mean (±SE) of the three replicate populations.

Copulation duration was significantly affected by Selection (*F* = 33.627, p < 0.001; Table S6A, Figure 1C), with Control populations having in general longer copulas than populations from the Warming regime. This trait was also affected by an interaction between Treatment and Mating Opportunity (*F* = 54.228, p < 0.001; Table S6A). Indeed, stressed males had shorter copulas than non-stressed males in the first mating opportunity; however, that pattern reversed from the second mating onward, with stressed males having longer copulation durations with increasing mating opportunities, and non-stressed males copulating faster in the second and third matings, with no differences between these last two (Table S7, Figure 1C). As in courtship and copulation latencies, History had no significant impact on copulation duration.

#### Effect of heatwave on male fertility

Male reproductive success was significantly shaped by the interaction between Treatment and Mating Opportunity in both the analysis of the Dutch and of the Portuguese populations (*X^2^* = 7.755, p = 0.021 for Dutch populations; *X^2^* = 7.996, p = 0.019 for Portuguese populations; Table S6B), but not by Selection. In both populations, this significant interaction resulted from stressed and non-stressed males displaying an opposite dynamic across mating opportunities, with stressed males having a slight increase in reproductive success in the second mating, contrary to non-stressed males whose reproductive success slightly decreases in that same mating (Table S8, Figure 2). Despite these different temporal dynamics associated with treatments, it is important to point out that no recovery of reproductive success occurred, stressed males continued to sire on average 56.6% fewer offspring than non-stressed males even in the last mating opportunity (Table S8, Figure 2; consistent patterns were observed for fecundity and offspring viability, Supplement 1).

**Figure 2.**
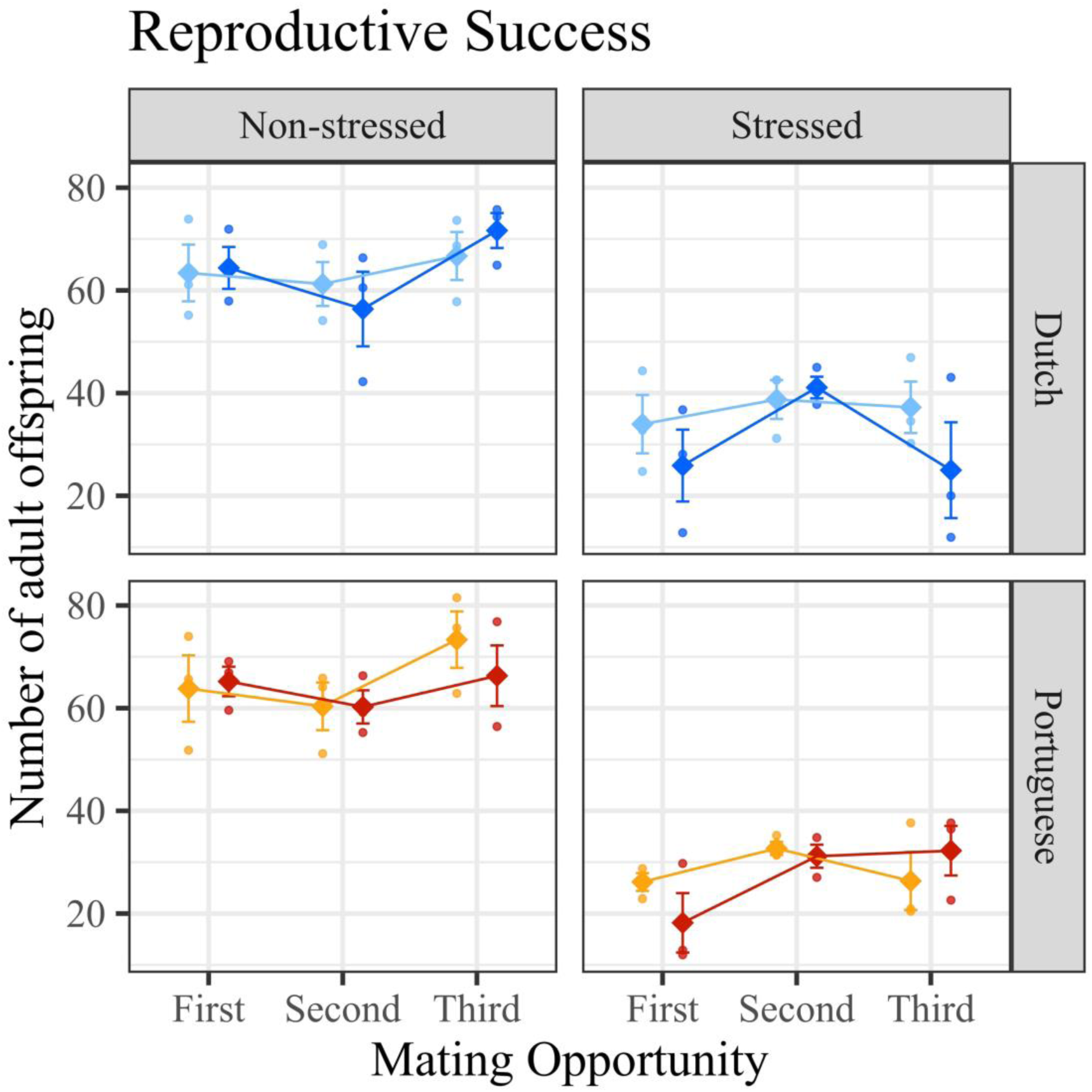
– Effect of heatwave during male adulthood on male fertility. Reproductive success: number of adult offspring in each vial. Colder colours represent Dutch populations, while warmer ones represent Portuguese populations. Lighter tones represent the Control regime, while darker tones represent the Warming regime. The small circles represent the mean values of each replicate population, and the big diamonds represent the mean (+SE) of the three replicate populations.

### Female mating behaviour and fertility following male heatwave

#### Effect of reduced male fertility on female mating behaviour

Courtship and copulation latencies exhibited identical responses, being significantly affected by the interaction between Treatment and Mating Opportunity (*X^2^* = 175.654, p < 0.001; *X^2^* = 207.797, p < 0.001, respectively; Table S9A, Figure 3A and 3B) but not by Selection or History. Indeed, when the first mating was with a stressed male, both courtship and copulation latency were much higher than when females first mated with a non-stressed male (see Table S10, Figure 3A and 3B). In addition, when the female first mated with a non-stressed male, the latency to court and copulate more than doubled in the second mating opportunity (see Table S10). A similar pattern was observed when females were paired twice with stressed males, with courtship and copulation latencies also increasing between the first and the second mating (Table S10) but showing higher absolute values than the control treatment. In turn, when females were first paired with a stressed male and then with a non-stressed male, both the courtship and copulation happened quicker in the second mating when compared to the first one (see Table S10, Figure 3A and 3B, respectively).

**Figure 3.**
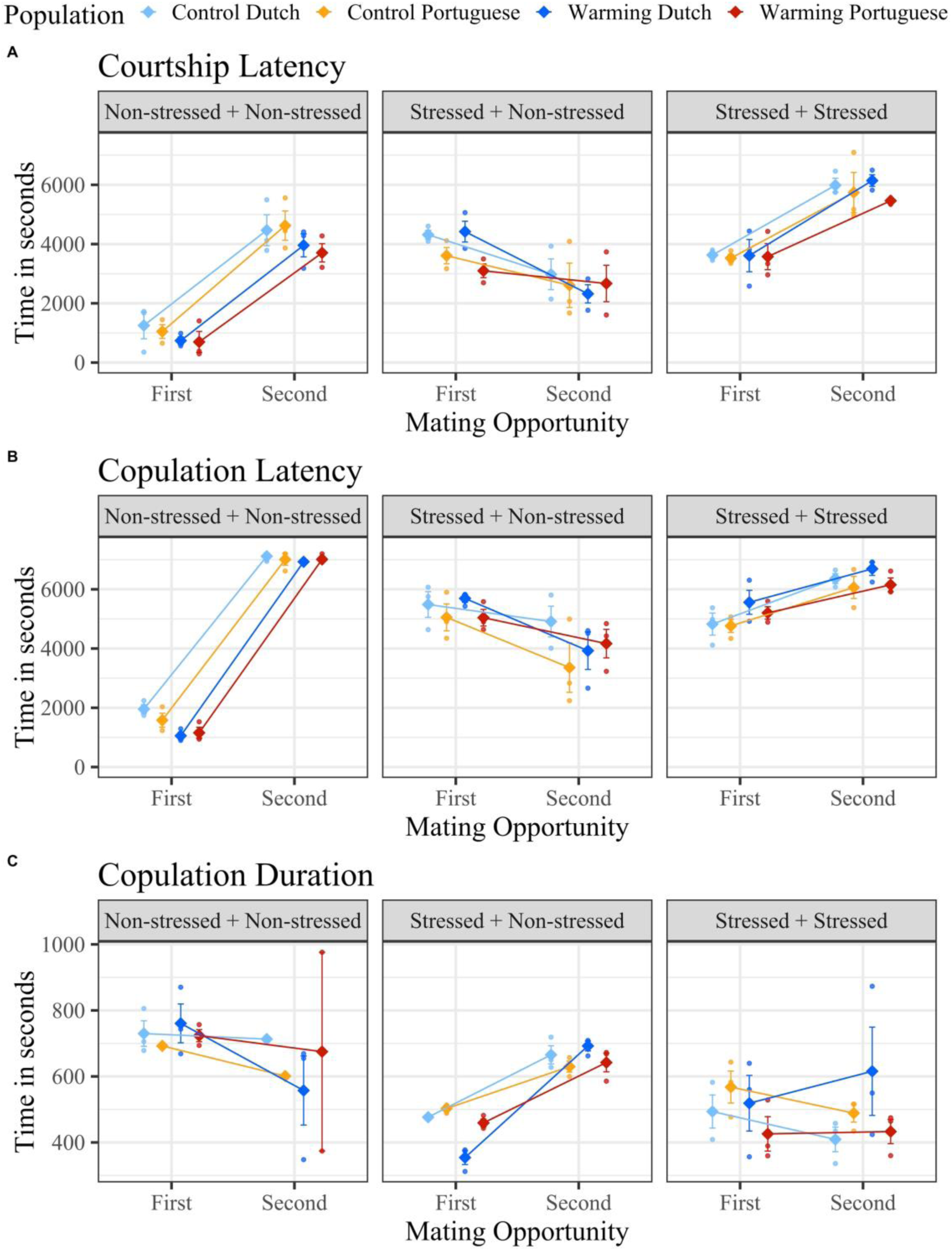
– Effect of heatwave during male adulthood on female mating behaviour. A) Courtship Latency: Time elapsed between pairing and beginning of male courtship; B) Copulation latency: Time elapsed between pairing and copulation beginning; C) Copulation Duration: Time elapsed between the beginning of the copula and its ending. Colder colours represent Dutch populations, while warmer ones represent Portuguese populations. Lighter tones represent the Control regime, while darker tones represent the Warming regime. The small circles represent the mean values of each replicate population, and the big diamonds represent the mean (+SE) of the three replicate populations.

Copulation duration was also significantly impacted by the interaction between Treatment and Mating Opportunity (F = 12.180, p < 0.001; see Table S9A, Figure 3C) and not by History or Selection. Indeed, copulation lasted longer when females mated with non-stressed males than with stressed males (see Table S10, Figure 3C), and the duration of matings with males from the same thermal treatment did not vary, irrespective of which combination of males the female was paired to, and of whether it was the female’s first or second mating opportunity.

#### Effect of reduced male fertility on female fertility

The reproductive success of both Dutch and Portuguese populations was significantly affected by the interaction between Treatment and Mating Opportunity (*X^2^* = 10.351, p = 0.006; *X^2^* = 14.701, p < 0.001, respectively; see Table S9B, Figure 4). In the first mating opportunity, both Dutch and Portuguese females that mated with stressed males had lower reproductive success than females paired with a non-stressed male (see Table S11A, Figure 4). Yet, from the first to the second mating opportunity, there was an overall increase of reproductive success, that differed depending on Treatment and History. Dutch females that were first paired with stressed and then with non-stressed males showed similar levels of reproductive success in the second mating opportunity relative to females that were paired with two non-stressed males (Table S11A). On the other hand, Dutch females paired with two stressed males approached, but did not reach, the levels of reproductive success of females that were paired with non-stressed males (Table S11A, Figure 4). In the Portuguese populations, the reproductive success of females that first mated with a stressed male, regardless of the thermal treatment of the second male, remained lower than that of females that first mated with non-stressed males (Table S11A, Figure 4). In addition, the reproductive success of Portuguese populations was also shaped by an interaction between Selection and Mating Opportunity (*X^2^* = 8.209, p = 0.004; see Table S9B), with females from the Control regime having lower reproductive success in the first mating opportunity than females from the Warming regime regardless of the treatment, but this difference disappeared in the second mating opportunity (Table S11B, Figure 4).

**Figure 4.**
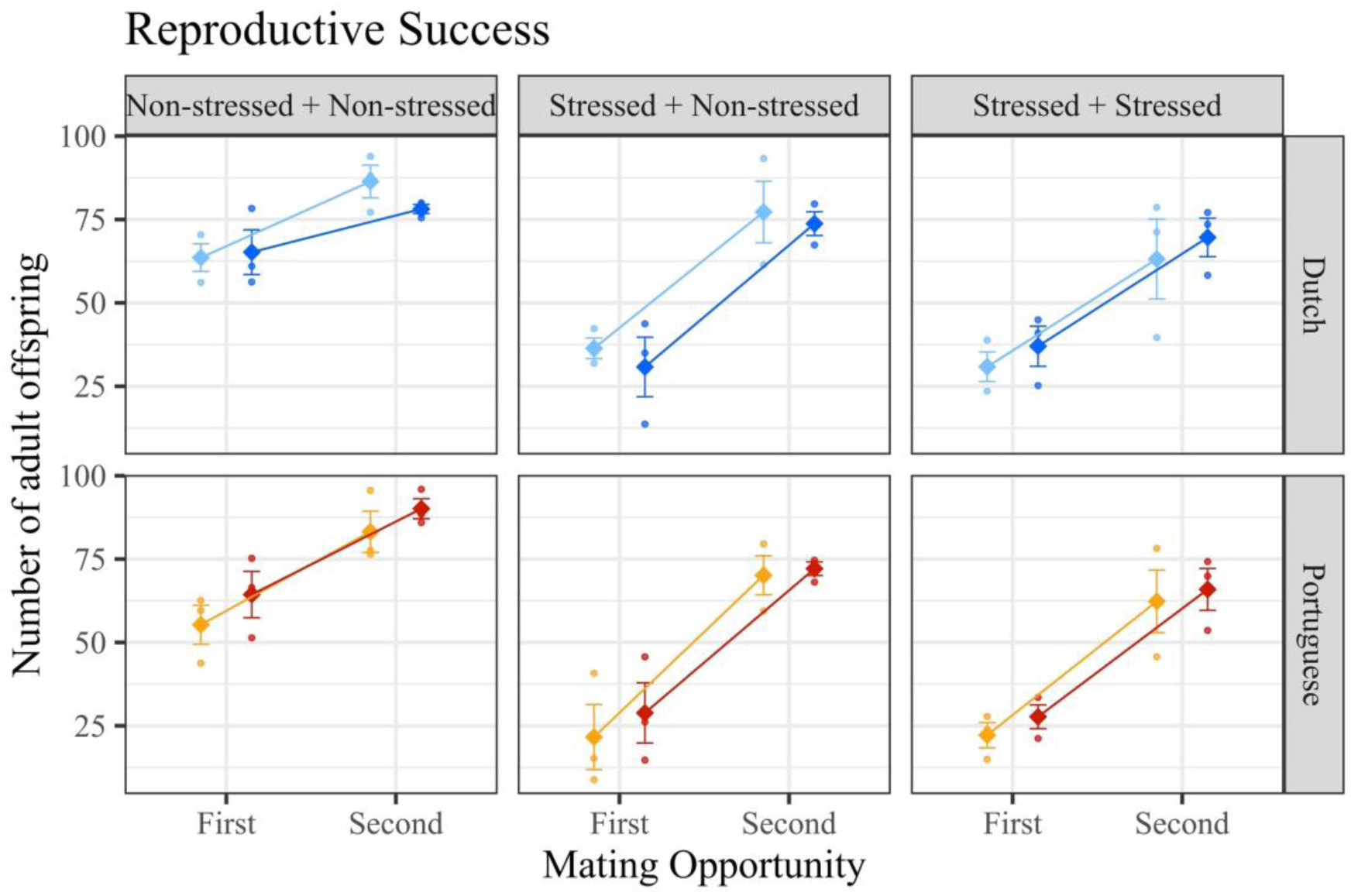
– Effect of heatwave during male adulthood on female fertility. Reproductive success: number of adult offspring in each vial. Colder colours represent Dutch populations, while warmer ones represent Portuguese populations. Lighter tones represent the Control regime, while darker tones represent the Warming regime. The small circles represent the mean values of each replicate population, and the big diamonds represent the mean (+SE) of the three replicate populations.

#### Effect of reduce male fertility on female propensity to remate & effect of increased remating rate on female reproductive success

Unlike females that first mated with a non-stressed male, females first mated with a male exposed to heat remated frequently (Treatment effect: *X^2^* = 77.706, p < 0.001; Table S12): remating frequency increased from 3% in females that first mated with a non-stressed male to 30% or more in females paired first with stressed males (30% in mated females paired with a second stressed male and 42% in mated females paired with a second non-stressed male; Table S13). No effect of History or Selection was found (Table S12).

As such, we set out to test whether the increase in remating was responsible (at least in part) for differences in reproductive success. The reproductive success of females that were paired with two non-stressed males was significantly and independently affected by both Mating Opportunity and Remating (*X^2^* = 100.527, p < 0.001; *X^2^* = 4.808, p = 0.028, respectively; see Table S14), with first matings resulting in lower reproductive success relative to the second ones and females that remated producing fewer offspring than those that did not (Figure 5, Table S15). Instead, when females were first paired with stressed males, their reproductive success was significantly shaped by an interaction between Mating Opportunity and Remating (*X^2^* = 40.931, p < 0.001, for the Stressed + non-stressed treatment; *X^2^* = 45.504, p < 0.001, for the Stressed + stressed treatment; Table S14). Indeed, in these cases, remated females showed a steeper increase in reproductive success across mating opportunities relative to non-remated females, such that the number of offspring produced after the second mating reached levels close to those found in non-remated females, even though in the first mating their reproductive success was lower (Figure 5, Table S15).

**Figure 5.**
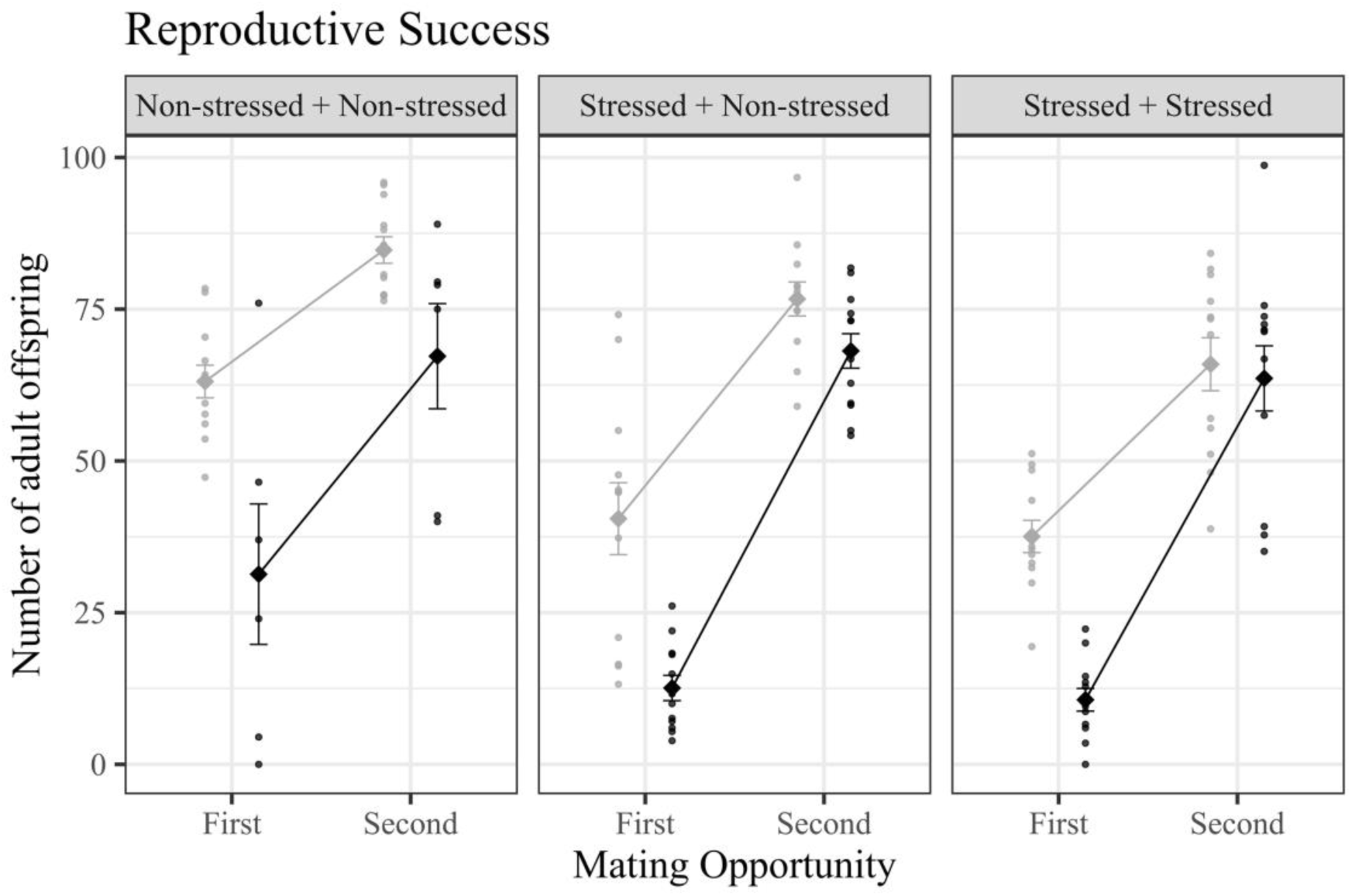
– Effect of remating behaviour on female fertility. Reproductive success: number of adult offspring in each vial. The colour grey represents females that did not remate, while the black colour represents remated females. The small circles represent the mean values of each replicate population, and the big diamonds represent the mean (±SE) of the twelve replicate populations.

## Discussion

### Male mating behaviour but not fertility recovers following adult heat stress

We observed a clear detrimental impact of high temperature during early adult stage on male mating behaviour in *Drosophila subobscura*. Specifically, stressed males took longer to start courting and mating after heat stress exposure and mated for a shorter period. Similar patterns have been reported in several studies in arthropods (Canal Domenech and Fricke, 2022; Costa et al., 2023; Leith et al., 2020; Mak et al., 2023; Sutter et al., 2019). This might be linked to a poor condition of heat-stressed males and/or to a lower female eagerness to accept males and mate for longer periods. Indeed, in *Drosophila subobscura* the beginning of courtship is mainly male-driven (Immonen et al., 2009), which suggests that heat-stressed males, who exhibited a longer latency to courtship, are less eager/able to mate. On the other hand, females from our Control populations more often accept nuptial feeding from non-stressed males than from males subjected to a heatwave (Grandela et al., 2024), which hints at the existence of female avoidance of stressed males. Curiously, a recent meta-analysis did not find clear evidence for the impact of temperature on mating latency and choosiness (Pilakouta and Baillet, 2022), which probably can be attributed to the lack of a consistent effect of temperature on mating behaviour due to differences in methodologies and/or in the species under study.

Interestingly, the immediate toll of heat on male’s mating behaviour was followed by recovery after the first mating in males from all selection regimes. Such a transversal immediate response points to a plastic rather than an evolutionary response. More specifically, there was a progressive decrease in latency to courtship and copula post-stress, with *D. subobscura* males recovering a significant portion of their behavioural performance within roughly a week after thermal stress (ca. 80% of recovery in courtship latency and 70% in copulation latency). Furthermore, copulation duration displayed three and eight days after exposure to heat stress increased, surpassing the duration of copulas with non-stressed males. Longer copulation duration was also found in *Tribolium castaneum* following thermal stress (Sales et al., 2018) and in *D. subobscura* under other stressful scenarios (Lizé et al., 2012). Less fertile males may benefit from extending their copula, if that allows them to maximise their reproductive output, especially considering increased female avoidance and assuming that the quality and/or the amount of functional sperm in stressed males is lower than in non-stressed males (e.g., Canal Domenech and Fricke, 2022; Sales et al., 2018). This might be particularly useful in a monandrous species such as *D. subobscura* whose females exit the mating pool after their first mating.

Exposure to sub-lethal temperatures, not only affected mating behaviour, but also led to a decline in male fertility, as reported in other studies (Costa et al., 2024; Parratt et al., 2021; Rodrigues et al., 2022; Sales et al., 2018; Walsh et al., 2021, 2019). However, unlike what was observed for behavioural traits, the fertility loss was permanent or at least long-lasting, with males remaining partially sterile a minimum of eight days post-stress. Evidence for recovery in arthropod males is equivocal: some studies find permanent sterility upon adult thermal stress (Meena et al., 2024; Parratt et al., 2021; Walsh et al., 2021), while in others, recovery is observed following heat exposure in either the developmental (Canal Domenech and Fricke, 2023, 2022; Sales et al., 2021; Walsh et al., 2021) or adult stages (Canal Domenech and Fricke, 2023; Parrett et al., 2024; Sales et al., 2021). Importantly, the lack of recovery in fertility that we observed after 8 days post-stress should have a high negative impact on male reproductive output, given the short life span of this species, particularly in warmer seasons (mostly less than a week, (Rosewell and Shorrocks, 1987). Still, the exposure to heat stress did not lead to full sterility, leaving room for the improvement in mating behaviour to confer some advantage to heat-stressed males by allowing them to (partially) fertilize more females. In this scenario, the recovery of mating behaviour (and the extension of mating duration) could be the only mechanism that heat-stressed males have to maximize progeny production after exposure to high temperatures and consequently prevent large population declines under global warming, a hypothesis that requires further testing.

### Increased female remating can rescue fertility when male sterility is prevalent

We showed that *D. subobscura* females are 10 to 14 times more prone to remate after mating with heat-stressed males. No effect of selection was observed in our results, suggesting that females plastically adjust their remating behaviour following mating with a stressed male. Similar results were reported in two studies with polyandrous species (2-fold increase in female remating for Drosophila pseudoobscura in Sutter et al., 2019; and around 1.3-fold increase for T. castaneum in Vasudeva et al., 2021). Several pre, peri and post-copulatory factors can help explain such a plastic response in remating behaviour. First, as discussed above, impaired male mating behaviour observed during first matings can act as a cue that elicits females to change their behaviour. Furthermore, there can be several factors that reduce the capacity of males to inhibit female remating, such as the disruption of male sex pheromones by temperature (Savarit and Ferveur, 2002) responsible for inhibiting female post-copulatory behaviour (Everaerts et al., 2010), the reduced transfer of functional seminal fluid proteins associated with diminished accessory gland size (Canal Domenech and Fricke, 2022), and a decrease in the levels of ejaculate stored in females’ spermatheca (Proshold, 1995). Future work should explore the relevance of these different mechanisms in explaining female remating behaviour and whether they vary across species.

The observed increase in remating resulted in improved female fertility, independently of the thermal treatment of the second male they were exposed to, indicating that females can boost their reproductive output by both remating with a non-stressed or heat-stressed male. In the case of females from Dutch populations, there was even a full restoration of fertility when they remated with non-stressed males, suggesting that population history may play a significant role in the females’ capacity to rescue their fertility and consequently in the population’s ability to withstand a heatwave. Still, studies with more populations from different latitudes are necessary to confirm this hypothesis. Importantly, and regardless of the level of the reproductive rescue, by disentangling the response of females that remated from that of females that do not, we were able to demonstrate that the remating behaviour, responsible for a steep increase in offspring production, is driving female reproductive rescue.

Our finding that *D. subobscura* females are strikingly more willing to remate after mating with a male exposed to a heatwave, is particularly enthralling given this species mating system. Indeed, because *D. subobscura* is monandrous (Fisher et al., 2013), the reported substantial increase in remating rates corresponds to a change of mating system – from monandry to polyandry. This creates an opportunity to better understand the complex eco-evolutionary consequences expected from a shift from single to multiple mating (Arnqvist and Nilsson, 2000; Holman and Kokko, 2013; Lizé et al., 2012; Moiron et al., 2022). Here we showed that this shift has consequences at the individual level, resulting from alterations in life-history traits related to fitness, namely in female reproductive output. However, many other changes are expected, namely in male-male and male-female interactions. For instance, our results suggest that under warming, females start using sperm from more than one male to produce progeny, creating the conditions for sperm competition to take place in *D. subobscura*. Likewise, with multiple paternity one might expect an increase in the genetic variability of progeny (Shuster et al., 2013), which may have important evolutionary implications, including on the likelihood of population persistence under global warming.

### Thermal selection did not enhance the response to a heatwave imposed on males

We have previously found an adaptive response to warming conditions in Portuguese *D. subobscura* populations, after 39 generations of thermal evolution (Santos et al., 2023), by assessing the reproductive success of warming and control populations in both (control or warming) environments. However, here, after 45 generations of thermal evolution, there was no strong evidence of evolutionary changes in mating behaviour or fertility of both males and females in response to a heatwave imposed on males. The only observed evolutionary responses found were for copulation duration, with males that evolved in a global warming scenario having shorter copulas, and for reproductive success, with Portuguese females that evolved under warming conditions producing more offspring than females evolved at benign temperatures but only in the first mating opportunity. Both responses could be related to the evolution of increased metabolic rate under warming conditions (Somero, 2012), as higher temperatures have been correlated to higher metabolic rates and higher activity in insects (Colinet et al., 2015; Tüzün and Stoks, 2022). In line with this, the populations subjected to the warming regime had a reduction of 3 days in developmental time when compared to populations that evolved at 18°C. All in all, rising temperatures seem to be linked to a fast-paced life (Tüzün and Stoks, 2022), resulting in quicker matings, and enhanced early fertility. It is nevertheless worth mentioning that the effect of thermal selection did not lead to overall increased offspring production, posing concerns about the efficacy of thermal adaptation in coping with sudden heatwave events.

Different explanations can be put forth for the lack of a strong evolutionary response in the present study. First, males were exposed to a thermal stress different from that experienced during thermal evolution. This mismatch reduces the possibility of detecting an evolutionary response and, more importantly, the lack of an adaptive response to the imposed heatwave suggests that adapting to certain warming conditions does not necessarily mean an advantage against warming in general terms. It could also be that the adaptive response hinges on male-female co-evolved traits. If that is the case, having used an outgroup as mates in our assays may have prevented the necessary male-female interactions to occur. Finally, it could be that an evolutionary response was hindered by the existing plastic response in female remating behaviour. This suggests that instead of favouring polyandrous females, selection may favour females that flexibly adjust their remating behaviour depending on environmental conditions (Gowaty, 2013). In the wild, where environmental variability is prevalent, this flexibility in remating behaviour may be crucial for population maintenance since the costs and benefits of polyandry are presumably changing dynamically (Holman and Kokko, 2013).

### Conclusions

All in all, our results show that both *D. subobscura* males and females can, to some extent, respond to the negative effects of exposure to sub-lethal temperatures. Interestingly, the extent of these responses was for the most part independent of the populations historical background and thermal selection. Instead, our results indicate that females from monandrous species, much like polyandrous ones, can make dynamic reproductive decisions, varying their mating behaviour according to their environment. Importantly, in monandrous species the observed plastic response in females corresponds to a plastic shift of mating system, from monandry to polyandry, a phenomenon that should not only buffer against the negative consequences of temperature but also lead to drastic eco-evolutionary rearrangements. The consequences thereof in the context of global warming are surprisingly understudied and merit further research, especially considering the increasing occurrence of heatwave events (IPCC 2022) and the recently found relevance of male thermal fertility limits in predicting the response of populations to global warming (Parratt et al., 2021; Walsh et al., 2019).

