## Supplementary material for "Monandrous flies do remate: plastic and evolutionary consequences of heat exposure on mating behaviour and fertility in *Drosophila subobscura*": Supplemetary Material

2 - Universidade de Lisboa, Lisbon, Portugal. Departamento de Biologia Animal, Faculdade de Ciências.

3 - Egas Moniz Center for Interdisciplinary Research (CiiEM), Egas Moniz School of Health and Science, Caparica, Portugal.

* - Co-last authors.

### Supplementary Material

**Table S1 – Description of the statistical models used for data analysis in *Recovery of male mating behaviour and fertility following heatwave*.** "Response variable": How the variable of interest was coded in the model. "Dataset": Data used to perform the analysis; when needed, the original data was divided into the different levels of a fixed factor present in the model: [a] – subset by History (Dutch or Portuguese). "Sample size": total number of replicates included in each analysis. [b] - Number of individuals per Mating Opportunity (First/Second/Third). "Maximal model": complete set of explanatory variables included in the model; random variables are represented within brackets. "Minimal model": model containing only the statistically significant variables. R function: The function used to code each model; square brackets indicate the error structure used ("g": gaussian; "qp1": quasi-Poisson, accounting for zero inflation; "p1": Poisson, accounting for zero inflation). "History": different bio-geographical origins of the populations under study (Dutch or Portuguese); "Selection": different thermal selective regimes of the populations under study (Control or Warming); "Treatment": different thermal treatments applied to the males under study (Non-stressed or Stressed); "Mating Opportunity": different mating events (First, Second or Third); "Block": different sets of same-numbered replicate populations (1, 2, or 3); "id": unique identification of each male. Courtship Latency: Time elapsed between pairing and beginning of male courtship; Copulation latency: Time elapsed between pairing and copulation beginning; Copulation Duration: Time elapsed between the beginning of the copula and its ending. Reproductive success: number of adult offspring in each vial. Censor: for both courtship and copulation latencies, whenever copulation was not observed within the time established for the mating assay, the data was censored.

| Var. of interest | Response variable | Dataset | Sample size | Maximal model | Minimal model | R function [err struct.] |
|---|---|---|---|---|---|---|
| Courtship latency | Surv(Courtship latency, censor) | All | 516/480/48 [b] | History * Selection * Treatment * Mating Opportunity + (1\|Block) | History + Selection + Treatment + Mating Opportunity + (Treatment:Mating Opportunity) + (1\|Block) | coxme [g] |
| Copulation latency | Surv(Copulation latency, censor) | All | 563/543/534 [b] | History * Selection * Treatment * Mating Opportunity + (1\|Block) | History + Selection + Treatment + Mating Opportunity + (Treatment:Mating Opportunity) + (1\|Block) | coxme [g] |
| Copulation duration | Copulation duration | All | 279/474/502 [b] | History * Selection * Treatment * Mating Opportunity + (1\|Block) | History + Selection + Treatment + Mating Opportunity + (Treatment:Mating Opportunity) + (1\|Block) | lmer [g] |
| Reproductive Success | Number of adult offspring | All | 562/542/533 [b] | History * Selection * Treatment * Mating Opportunity + (1\|Block) + (Mating Opportunity\|id) | History * Selection * Treatment * Mating Opportunity + (1\|Block) + (Mating Opportunity\|id) | glmmTMB [qp1] |
| | | Dutch [a] | 277/263/259 [b] | Selection * Treatment * Mating Opportunity + (1\|Block) + (Mating Opportunity\|id) | Selection + Treatment + Mating Opportunity + (Treatment:Mating Opportunity) + (1\|Block) + (Mating Opportunity\|id) | glmmTMB [p1] |
| | | Portuguese [a] | 285/279/274 [b] | Selection * Treatment * Mating Opportunity + (1\|Block) + (Mating Opportunity\|id) | Selection + Treatment + Mating Opportunity + (Treatment:Mating Opportunity) + (1\|Block) + (Mating Opportunity\|id) | glmmTMB [qp1] |

**Table S2 – Description of the statistical models used for data analysis in *Female mating behaviour and fertility following male heatwave*.** "Response variable": How the variable of interest was coded in the model. "Dataset": Data used to perform the analysis; when needed, the original data was divided into the different levels of a fixed factor present in the model: [a] – subset by History (Dutch or Portuguese). [b] – subset by Treatment (Non-stressed or Stressed). "Sample size": total number of replicates included in each analysis. [c] - number of individuals per Mating Opportunity (First/Second). "Maximal model": complete set of explanatory variables included in the model; random variables are represented within brackets. "Minimal model": model containing only the statistically significant variables. R function: The function used to code each model; square brackets indicate the error structure used ("g": gaussian; "qp1": quasi-Poisson, accounting for zero inflation; "b": binomial; "nb1"; negative binomial, accounting for zero inflation "qp": quasi-Poisson). "History": different bio-geographical origins of the populations under study (Dutch or Portuguese); "Selection": different thermal selective regimes of the populations under study (Control or Warming); "Treatment": different combinations of males paired with the females under study (Non-stressed + non-stressed, Stressed + non-stressed or Stressed + stressed); "Mating Opportunity": different mating events (First or Second); "Block": different sets of same-numbered replicate populations (1, 2, or 3); "id": unique identification of each female; "Remating" difference between non-remated and remated females (No Remating or Remated). Remating: Female mates with a different male after already mating once. Courtship Latency: Time elapsed between pairing and beginning of male courtship; Copulation latency: Time elapsed between pairing and copulation beginning; Copulation Duration: Time elapsed between the beginning of the copula and its ending. Reproductive success: number of adult offspring in each vial. Censor: for both courtship and copulation latencies, whenever copulation was not observed within the time established for the mating assay, the data was censored. Note that to confirm that the remating behaviour was responsible for differences in reproductive success, an additional analysis of reproductive success with Remating as a fixed factor was performed.

| **Var. of interest** | **Response variable** | **Dataset** | **Sample size** | **Maximal model** | **Minimal model** | **R subroutine [err struct.]** |
|---|---|---|---|---|---|---|
| Remating | Remating | All | 834 | History * Selection * Treatment + (1\|Block) | History + Selection + Treatment + (1\|Block) | glmmTMB [b] |
| Courtship latency | Surv(Courtship latency, censor) | All | 795/794 [c] | History * Selection * Treatment * Mating Opportunity + (1\|Block) | History + Selection + Treatment + Mating Opportunity + (Treatment:Mating Opportunity) + (1\|Block) | coxme [g] |
| Copulation latency | Surv(Copulation latency, censor) | All | 849/833 [c] | History * Selection * Treatment*Mating Opportunity + (1\|Block) | History + Selection + Treatment + Mating Opportunity + (Treatment:Mating Opportunity) + (1\|Block) | coxme [g] |
| Copulation duration | Copulation duration | All | 489/191 [c] | History * Selection * Treatment * Mating Opportunity + (1\|Block) | History + Selection + Treatment + Mating Opportunity + (Treatment:Mating Opportunity) + (1\|Block) | lmer [g] |
| Reproductive Success | Number of adult offspring | All | 842/833 [c] | History * Selection * Treatment * Mating Opportunity + (1\|Block) + (Mating Opportunity\|id) | History + Selection + Treatment + Mating Opportunity + (Treatmentt:Mating Opportunity) + (1\|Block) + (Mating Opportunity\|id) | glmmTMB [qp1] |
| | | Dutch [a] | 419/415 [c] | Selection * Treatment * Mating Opportunity + (1\|Block) + (Mating Opportunity\|id) | Selection + Treatment + Mating Opportunity + (Treatment:Mating Opportunity )+ (1\|Block) + (Mating Opportunity\|id) | glmmTMB [qp1] |
| | | Portuguese [a] | 423/418 [c] | Selection * Treatment * Mating Opportunity + (1\|Block) + (Mating Opportunity\|id) | Selection + Treatment + Mating Opportunity + (Selection:Mating Opportunity) + (Treatment:Mating Opportunity) + (1\|Block)+(Mating Opportunity\|id) | glmmTMB [qp1] |
| | | All | 842/833 [c] | Treatment * Mating Opportunity * Remating + (1\|Block) + (Mating Opportunity\|id) | Treatment * Mating Opportunity * Remating + (1\|Block) + (Mating Opportunity\|id) | glmmTMB [qp1] |
| | | Non-stressed + non-stressed [b] | 286/284 [c] | Mating Opportunity * Remating + (1\|Block) + (Mating Opportunity\|id) | Mating Opportunity + Remating + (1\|Block) + (Mating Opportunity\|id) | glmmTMB [nb1] |
| | | Stressed + non-stressed [b] | 287/285 [c] | Mating Opportunity * Remating + (1\|Block) + (Mating Opportunity\|id) | Mating Opportunity * Remating + (1\|Block) + (Mating Opportunity\|id) | glmmTMB [qp] |
| | | Stressed + stressed [b] | 269/264 [c] | Mating Opportunity * Remating + (1\|Block) + (Mating Opportunity\|id) | Mating Opportunity * Remating + (1\|Block) + (Mating Opportunity\|id) | glmmTMB [qp1] |

**Table S3 – Results from the analyses of variances of the effect of heatwave during male adulthood on male fertility.** Reproductive success: number of adult offspring in each vial. Males with different bio-geographical origins ("History": Dutch or Portuguese) that were exposed for 45 generations to different thermal conditions ("Selection": Control or Warming), were subjected, or not, to a heatwave treatment ("Treatment": Non-stressed or Stressed) and were paired, with different females, 1, 3 and 8 days after the thermal treatment ("Mating Opportunity": First, Second or Third). "Df": the degrees of freedom. "$X^2$": the Chi-square value obtained in each analysis. Statistically significant terms are represented in bold.

| Trait | Independent Variable | *Df* | $X^2$ | *p-value* |
|---|---|---|---|---|
| Reproductive Success | History | 1 | 0 | 0.997 |
| | Selection | 1 | 0.243 | 0.622 |
| | **Treatment** | **1** | **105.811** | **< 0.001** |
| | Mating Opportunity | 2 | 2.381 | 0.304 |
| | History*Selection | 1 | 0.024 | 0.877 |
| | History*Treatment | 1 | 0.080 | 0.778 |
| | Selection*Treatment | 1 | 0.193 | 0.661 |
| | History*Mating Opportunity | 2 | 2.871 | 0.237 |
| | Selection*Mating Opportunity | 2 | 2.367 | 0.306 |
| | **Treatment*Mating Opportunity** | **2** | **17.554** | **< 0.001** |
| | History*Selection*Treatment | 1 | 0.002 | 0.965 |
| | History*Selection*Mating Opportunity | 2 | 1.863 | 0.394 |
| | History*Treatment*Mating Opportunity | 2 | 2.366 | 0.306 |
| | Selection*Treatment*Mating Opportunity | 2 | 2.588 | 0.274 |
| | **History*Selection*Treatment*Mating Opportunity** | **2** | **7.441** | **0.024** |

**Table S4 – Results from the analyses of variances of the effect of heatwave during male adulthood on female fertility.** Reproductive success: number of adult offspring in each vial. Females with different bio-geographical origins ("History": Dutch or Portuguese), that were exposed for 45 generations to different thermal conditions ("Selection": Control or Warming), were paired twice with different males subjected, or not, to a heatwave treatment ("Treatment": Non-stressed + non-stressed, Stressed + non-stressed or Stressed + stressed), 1 and 3 days after the male thermal treatment ("Mating Opportunity": First or Second). "Df": the degrees of freedom. "$X^2$": the Chi-square value obtained in each analysis. Statistically significant terms are represented in bold.

| Trait | Explanatory Variable | *Df* | $X^2$ | *p-value* |
|---|---|---|---|---|
| Reproductive Success | **History** | **1** | **6.668** | **0.010** |
| | **Selection** | **1** | **5.196** | **0.023** |
| | **Treatment** | **2** | **97.506** | **< 0.001** |
| | **Mating Opportunity** | **1** | **335.746** | **< 0.001** |
| | **History*Selection** | **1** | **6.660** | **0.010** |
| | **History*Mating Opportunity** | **1** | **7.980** | **0.010** |
| | **Selection*Mating Opportunity** | **1** | **8.686** | **0.005** |
| | **Treatment*Mating Opportunity** | **2** | **22.884** | **< 0.001** |

**Table S5 – Results from the analyses of variances of the effect of remating behaviour on the female reproductive success.** Reproductive success: number of adult offspring in each vial. Females paired twice with different males subjected, or not, to a heatwave treatment ("Treatment": Non-stressed + non-stressed, Stressed + non-stressed or Stressed + stressed), 1 and 3 days after the male thermal treatment ("Mating Opportunity": First or Second), displayed remating, or not ("Remating": No Remating or Remated). "Df": the degrees of freedom. "$X^2$": the Chi-square value obtained in each analysis. Statistically significant terms are represented in bold.

| Trait | Explanatory Variable | *Df* | $X^2$ | *p-value* |
|---|---|---|---|---|
| Reproductive Success | **Treatment** | **2** | **59.732** | **< 0.001** |
| | **Mating Opportunity** | **1** | **271.638** | **< 0.001** |
| | **Remating** | **1** | **161.840** | **< 0.001** |
| | **Treatment*Mating Opportunity** | **2** | **42.999** | **< 0.001** |
| | **Treatment*Remating** | **2** | **30.161** | **< 0.001** |
| | **Mating Opportunity*Remating** | **1** | **124.169** | **< 0.001** |
| | **Treatment*Mating Opportunity*Remating** | **2** | **34.411** | **< 0.001** |

**Table S6 – Results from the analyses of variances of the effect of heatwave during male adulthood on male (A) mating behaviour and (B) fertility.** Courtship Latency: Time elapsed between pairing and beginning of male courtship; Copulation latency: Time elapsed between pairing and copulation beginning; Copulation Duration: Time elapsed between the beginning of the copula and its ending. Reproductive success: number of adult offspring in each vial (for this trait populations with different History were analysed separately). Males with different bio-geographical origins ("History": Dutch or Portuguese) that were exposed for 45 generations to different thermal conditions ("Selection": Control or Warming), were subjected, or not, to a heatwave treatment ("Treatment": Non-stressed or Stressed) and were paired, with different females, 1, 3 and 8 days after the thermal treatment ("Mating Opportunity": First, Second or Third). "Df": the degrees of freedom. Df.res: residual degrees of freedom. "$X^2$": the Chi-square value obtained in each analysis. "F": F-test obtained in each analysis. Statistically significant terms are represented in bold.

A)

| Trait | Independent Variable | *Df (Df.res)* | *Tests statistics* | *p-value* |
|---|---|---|---|---|
| | | | $X^2$ | |
| Courtship Latency | History | 1 | 0.537 | 0.464 |
| | Selection | 1 | 0.968 | 0.325 |
| | **Treatment** | **1** | **247.200** | **< 0.001** |
| | **Mating Opportunity** | **2** | **281.521** | **< 0.001** |
| | **Treatment*Mating Opportunity** | **2** | **37.469** | **< 0.001** |
| | | | $X^2$ | |
| Copulation Latency | History | 1 | 1.748 | 0.186 |
| | Selection | 1 | 0.028 | 0.868 |
| | **Treatment** | **1** | **359.925** | **< 0.001** |
| | **Mating Opportunity** | **2** | **290.249** | **< 0.001** |
| | **Treatment*Mating Opportunity** | **2** | **33.060** | **< 0.001** |
| | | | *F* | |
| Copulation Duration | History | 1 (1245.360) | 0.602 | 0.438 |
| | **Selection** | **1 (1245.060)** | **33.627** | **< 0.001** |
| | **Treatment** | **1 (1246.090)** | **69.773** | **< 0.001** |
| | **Mating Opportunity** | **2 (1245.950)** | **15.100** | **< 0.001** |
| | **Treatment*Mating Opportunity** | **2 (1245.900)** | **54.228** | **< 0.001** |

B)

| **History** | **Trait** | **Independent Variable** | ***Df*** | **$X^2$** | ***p-value*** |
|---|---|---|---|---|---|
| Dutch | Reproductive Success | Selection | 1 | 0.003 | 0.958 |
| | | **Treatment** | **1** | **47.181** | **< 0.001** |
| | | Mating Opportunity | 2 | 0.170 | 0.919 |
| | | **Treatment*Mating Opportunity** | **2** | **7.755** | **0.021** |
| Portuguese | Reproductive Success | Selection | 1 | 0.040 | 0.842 |
| | | **Treatment** | **1** | **51.609** | **< 0.001** |
| | | Mating Opportunity | 2 | 3.468 | 0.177 |
| | | **Treatment*Mating Opportunity** | **2** | **7.966** | **0.019** |

**Table S7 – *A posteriori* contrasts for the effect of heatwave during male adulthood on male mating behaviour.** Courtship Latency: Time elapsed between pairing and beginning of male courtship; Copulation latency: Time elapsed between pairing and copulation beginning; Copulation Duration: Time elapsed between the beginning of the copula and its ending. *A posteriori* tukey contrasts. "Z or T ratio": the T-test value obtained in each comparison. Comparison: Interaction between Treatment (Stressed or Non-stressed) and Mating Opportunity (First, Second or Third). Statistically significant terms are represented in bold.

| Trait | Comparison | *Z or T ratio* | *p-value* |
|---|---|---|---|
| | | *Z ratio* | |
| Courtship Latency | **First Non-stressed - First Stressed** | **13.427** | **<0.001** |
| | First Non-stressed - Second Stressed | 1.533 | 0.643 |
| | First Non-stressed - Third Stressed | -1.923 | 0.388 |
| | **First Non-stressed - Second Non-stressed** | **-7.778** | **<0.001** |
| | **First Non-stressed - Third Non-stressed** | **-7.297** | **<0.001** |
| | **Second Non-stressed - First Stressed** | **-20.018** | **<0.001** |
| | **Second Non-stressed - Second Stressed** | **9.096** | **<0.001** |
| | **Second Non-stressed - Third Stressed** | **5.816** | **<0.001** |
| | Second Non-stressed - Third Non-stressed | 0.465 | 0.997 |
| | **Third Non-stressed - First Stressed** | **-19.424** | **<0.001** |
| | **Third Non-stressed - Second Stressed** | **-8.615** | **<0.001** |
| | **Third Non-stressed - Third Stressed** | **5.357** | **<0.001** |
| | **First Stressed - Second Stressed** | **-11.906** | **<0.001** |
| | **First Stressed - Third Stressed** | **-14.894** | **<0.001** |
| | **Second Stressed - Third Stressed** | **-3.403** | **0.009** |
| | | *Z ratio* | |
| Copulation Latency | **First Non-stressed - First Stressed** | **14.902** | **<0.001** |
| | **First Non-stressed - Second Stressed** | **3.635** | **0.004** |
| | First Non-stressed - Third Stressed | -0.284 | 0.999 |
| | **First Non-stressed - Second Non-stressed** | **-8.206** | **<0.001** |
| | **First Non-stressed - Third Non-stressed** | **-7.899** | **<0.001** |
| | **Second Non-stressed - First Stressed** | **-21.717** | **<0.001** |
| | **Second Non-stressed - Second Stressed** | **11.453** | **<0.001** |
| | **Second Non-stressed - Third Stressed** | **7.684** | **<0.001** |
| | Second Non-stressed - Third Non-stressed | 0.210 | 0.999 |
| | **Third Non-stressed - First Stressed** | **-21.142** | **<0.001** |
| | **Third Non-stressed - Second Stressed** | **-11.114** | **<0.001** |
| | **Third Non-stressed - Third Stressed** | **7.408** | **<0.001** |
| | **First Stressed - Second Stressed** | **-11.292** | **<0.001** |
| | **First Stressed - Third Stressed** | **-14.729** | **<0.001** |
| | **Second Stressed - Third Stressed** | **3.802** | **0.002** |
| | | *T ratio* | |
| Copulation Duration | **First Non-stressed - First Stressed** | **3.816** | **0.002** |
| | **First Non-stressed - Second Stressed** | **-5.902** | **<0.001** |
| | **First Non-stressed - Third Stressed** | **-9.690** | **<0.001** |
| | **First Non-stressed - Second Non-stressed** | **6.642** | **<0.001** |
| | **First Non-stressed - Third Non-stressed** | **5.595** | **<0.001** |
| | Second Non-stressed - First Stressed | -0.555 | 0.994 |
| | **Second Non-stressed - Second Stressed** | **-12.367** | **<0.001** |
| | **Second Non-stressed - Third Stressed** | **-16.485** | **<0.001** |
| | Second Non-stressed - Third Non-stressed | -1.083 | 0.888 |
| | Third Non-stressed - First Stressed | -1.075 | 0.891 |
| | **Third Non-stressed - Second Stressed** | **11.375** | **<0.001** |
| | **Third Non-stressed - Third Stressed** | **-15.464** | **<0.001** |
| | **First Stressed - Second Stressed** | **-6.852** | **<0.001** |
| | **First Stressed - Third Stressed** | **-8.746** | **<0.001** |
| | **Second Stressed - Third Stressed** | **-3.433** | **0.008** |

**Table S8 – *A posteriori* contrasts for the effect of heatwave during male adulthood on male fertility for Dutch and Portuguese populations separately.** Reproductive success: number of adult offspring in each vial. *A posteriori* tukey contrasts. “T ratio”: the T-test value obtained in each comparison. Comparison: Interaction between Treatment (Stressed or Non-stressed) and Mating Opportunity (First, Second or Third). Statistically significant terms are represented in bold.

| History | Trait | Comparison | *T ratio* | *p-value* |
|---|---|---|---|---|
| Dutch | Reproductive Success | **First Non-stressed - First Stressed** | **5.109** | **<0.001** |
| | | **First Non-stressed - Second Stressed** | **3.919** | **0.001** |
| | | **First Non-stressed - Third Stressed** | **5.759** | **<0.001** |
| | | First Non-stressed - Second Non-stressed | 1.869 | 0.422 |
| | | First Non-stressed - Third Non-stressed | -0.737 | 0.977 |
| | | **Second Non-stressed - First Stressed** | **-3.370** | **0.010** |
| | | Second Non-stressed - Second Stressed | 2.309 | 0.192 |
| | | **Second Non-stressed - Third Stressed** | **3.989** | **0.001** |
| | | Second Non-stressed - Third Non-stressed | -2.447 | 0.142 |
| | | **Third Non-stressed - First Stressed** | **-5.662** | **<0.001** |
| | | **Third Non-stressed - Second Stressed** | **-4.456** | **<0.001** |
| | | **Third Non-stressed - Third Stressed** | **6.321** | **<0.001** |
| | | First Stressed - Second Stressed | -0.969 | 0.928 |
| | | First Stressed - Third Stressed | 0.636 | 0.988 |
| | | Second Stressed - Third Stressed | 1.547 | 0.634 |
| Portuguese | Reproductive Success | **First Non-stressed - First Stressed** | **5.745** | **<0.001** |
| | | **First Non-stressed - Second Stressed** | **3.616** | **0.004** |
| | | **First Non-stressed - Third Stressed** | **4.201** | **<0.001** |
| | | First Non-stressed - Second Non-stressed | 1.388 | 0.735 |
| | | First Non-stressed - Third Non-stressed | -0.852 | 0.958 |
| | | **Second Non-stressed - First Stressed** | **-4.721** | **<0.001** |
| | | Second Non-stressed - Second Stressed | 2.499 | 0.125 |
| | | **Second Non-stressed - Third Stressed** | **3.118** | **0.023** |
| | | Second Non-stressed - Third Non-stressed | -2.323 | 0.186 |
| | | **Third Non-stressed - First Stressed** | **-6.303** | **<0.001** |
| | | **Third Non-stressed - Second Stressed** | **-4.267** | **<0.001** |
| | | **Third Non-stressed - Third Stressed** | **4.839** | **<0.001** |
| | | First Stressed - Second Stressed | -2.332 | 0.182 |
| | | First Stressed - Third Stressed | -1.651 | 0.565 |
| | | Second Stressed - Third Stressed | 0.678 | 0.984 |

**Table S9– Results from the analyses of variances of the effect of heatwave during male adulthood on female (A) mating behaviour and (B) fertility.** Courtship Latency: Time elapsed between pairing and beginning of male courtship; Copulation latency: Time elapsed between pairing and copulation beginning; Copulation Duration: Time elapsed between the beginning of the copula and its ending. Reproductive success: number of adult offspring in each vial (for this trait populations with different History were analysed separately). Females with different bio-geographical origins ("History": Dutch or Portuguese), that were exposed for 45 generations to different thermal conditions ("Selection": Control or Warming), were paired twice with different males subjected, or not, to a heatwave treatment ("Treatment": Non-stressed + non-stressed, Stressed + non-stressed or Stressed + stressed), 1 and 3 days after the male thermal treatment ("Mating Opportunity": First or Second). "Df": the degrees of freedom. Df.res: residual degrees of freedom. "$X^2$": the Chi-square value obtained in each analysis. "F": F-test obtained in each analysis. Statistically significant terms are represented in bold.

A)

| Trait | Independent Variable | *Df (Df.res)* | *Test statistics* | *p-value* |
|---|---|---|---|---|
| | | | ***$X^2$*** | |
| Courtship Latency | History | 1 | 3.150 | 0.076 |
| | Selection | 1 | 0.756 | 0.385 |
| | **Treatment** | **2** | **54.491** | **< 0.001** |
| | **Mating Opportunity** | **1** | **290.333** | **< 0.001** |
| | **Treatment*Mating Opportunity** | **2** | **175.654** | **< 0.001** |
| | | | ***$X^2$*** | |
| Copulation Latency | History | 1 | 2.539 | 0.111 |
| | Selection | 1 | 0.001 | 0.982 |
| | **Treatment** | **2** | **48.543** | **< 0.001** |
| | **Mating Opportunity** | **1** | **361.207** | **< 0.001** |
| | **Treatment*Mating Opportunity** | **2** | **207.797** | **< 0.001** |
| | | | ***F*** | |
| Copulation Duration | History | 1 (670.640) | 0.711 | 0.400 |
| | Selection | 1 (670.690) | 0.188 | 0.665 |
| | **Treatment** | **2 (670.450)** | **10.190** | **< 0.001** |
| | Mating Opportunity | 1 (671.210) | 1.288 | 0.257 |
| | **Treatment*Mating Opportunity** | **2 (670.340)** | **12.180** | **< 0.001** |

B)

| History | Trait | Explanatory Variable | *Df* | *$X^2$* | *p-value* |
|---|---|---|---|---|---|
| Dutch | Reproductive Success | Selection | 1 | 0.690 | 0.406 |
| | | **Treatment** | **2** | **35.879** | **< 0.001** |
| | | **Mating Opportunity** | **1** | **137.913** | **< 0.001** |
| | | **Treatment*Mating Opportunity** | **2** | **10.351** | **0.006** |
| Portuguese | Reproductive Success | **Selection** | **1** | **12.401** | **< 0.001** |
| | | **Treatment** | **2** | **63.761** | **< 0.001** |
| | | **Mating Opportunity** | **1** | 188.398 | **< 0.001** |
| | | **Selection*Mating Opportunity** | **1** | **8.209** | **0.004** |
| | | **Treatment*Mating Opportunity** | **2** | **14.701** | **< 0.001** |

**Table S10 – *A posteriori* contrasts for the effect of heatwave during male adulthood on female mating behaviour.** Courtship Latency: Time elapsed between pairing and beginning of male courtship; Copulation latency: Time elapsed between pairing and copulation beginning; Copulation Duration: Time elapsed between the beginning of the copula and its ending. *A posteriori* tukey contrasts. “Z or T ratio”: the T-test value obtained in each comparison. Comparison: Interaction between Treatment (Non-stressed + non-stressed, Stressed + non-stressed or Stressed + stressed) and Mating Opportunity (First or Second). Statistically significant terms are represented in bold.

| Trait | Comparison | *Z or T ratio* | *p-value* |
|---|---|---|---|
| | | **Z ratio** | |
| Courtship Latency | **First Non-stressed + non-stressed – First Stressed + non-stressed** | **12.519** | **< 0.001** |
| | **First Non-stressed + non-stressed – Second Stressed + non-stressed** | **13.798** | **< 0.001** |
| | **First Non-stressed + non-stressed – First Stressed + stressed** | **10.787** | **< 0.001** |
| | **First Non-stressed + non-stressed – Second Stressed + stressed** | **20.049** | **< 0.001** |
| | **First Non-stressed + non-stressed - Second Non-stressed + non-stressed** | **13.543** | **< 0.001** |
| | **Second Non-stressed + non-stressed – First Stressed + non-stressed** | **10.315** | **< 0.001** |
| | **Second Non-stressed + non-stressed – Second Stressed + non-stressed** | **-9.292** | **< 0.001** |
| | **Second Non-stressed + non-stressed – First Stressed + stressed** | **10.817** | **< 0.001** |
| | **Second Non-stressed + non-stressed – Second Stressed + stressed** | **-6.224** | **< 0.001** |
| | First Stressed + non-stressed – First Stressed + stressed | -1.961 | 0.365 |
| | **First Stressed + non-stressed – Second Stressed + stressed** | **11.203** | **< 0.001** |
| | **First Stressed + non-stressed – Second Stressed + non-stressed** | **3.079** | **0.025** |
| | **Second Stressed + non-stressed – First Stressed + stressed** | **4.767** | **< 0.001** |
| | **Second Stressed + non-stressed – Second Stressed + stressed** | **7.901** | **< 0.001** |
| | **First Stressed + stressed – Second Stressed + stressed** | **12.633** | **< 0.001** |
| | | **Z ratio** | |
| Copulation Latency | **First Non-stressed + non-stressed – First Stressed + non-stressed** | **14.245** | **< 0.001** |
| | **First Non-stressed + non-stressed – Second Stressed + non-stressed** | **16.352** | **< 0.001** |
| | **First Non-stressed + non-stressed – First Stressed + stressed** | **12.339** | **< 0.001** |
| | **First Non-stressed + non-stressed – Second Stressed + stressed** | **20.637** | **< 0.001** |
| | **First Non-stressed + non-stressed - Second Non-stressed + non-stressed** | **15.716** | **< 0.001** |
| | **Second Non-stressed + non-stressed – First Stressed + non-stressed** | **11.780** | **< 0.001** |
| | **Second Non-stressed + non-stressed – Second Stressed + non-stressed** | **-10.461** | **< 0.001** |
| | **Second Non-stressed + non-stressed – First Stressed + stressed** | **12.354** | **< 0.001** |
| | **Second Non-stressed + non-stressed – Second Stressed + stressed** | **-7.748** | **< 0.001** |
| | First Stressed + non-stressed – First Stressed + stressed | -2.091 | 0.292 |
| | **First Stressed + non-stressed – Second Stressed + stressed** | **10.264** | **< 0.001** |
| | **First Stressed + non-stressed – Second Stressed + non-stressed** | **3.925** | **0.001** |
| | **Second Stressed + non-stressed – First Stressed + stressed** | **5.764** | **< 0.001** |
| | **Second Stressed + non-stressed – Second Stressed + stressed** | **6.548** | **< 0.001** |
| | **First Stressed + stressed – Second Stressed + stressed** | **11.812** | **< 0.001** |
| | | **T ratio** | |
| Copulation Duration | **First Non-stressed + non-stressed – First Stressed + non-stressed** | **9.978** | **< 0.001** |
| | First Non-stressed + non-stressed – Second Stressed + non-stressed | 2.423 | 0.150 |
| | **First Non-stressed + non-stressed – First Stressed + stressed** | **9.408** | **< 0.001** |
| | **First Non-stressed + non-stressed – Second Stressed + stressed** | **6.439** | **< 0.001** |
| | First Non-stressed + non-stressed - Second Non-stressed + non-stressed | 1.123 | 0.872 |
| | Second Non-stressed + non-stressed – First Stressed + non-stressed | -2.037 | 0.322 |
| | Second Non-stressed + non-stressed – Second Stressed + non-stressed | -0.403 | 0.999 |
| | Second Non-stressed + non-stressed – First Stressed + stressed | -1.714 | 0.523 |
| | Second Non-stressed + non-stressed – Second Stressed + stressed | 1.665 | 0.556 |
| | First Stressed + non-stressed – First Stressed + stressed | -0.918 | 0.942 |
| | First Stressed + non-stressed – Second Stressed + stressed | -0.611 | 0.990 |
| | **First Stressed + non-stressed – Second Stressed + non-stressed** | **-6.891** | **< 0.001** |
| | **Second Stressed + non-stressed – First Stressed + stressed** | **-6.188** | **< 0.001** |
| | **Second Stressed + non-stressed – Second Stressed + stressed** | **4.579** | **0.001** |
| | First Stressed + stressed – Second Stressed + stressed | 0.072 | 1 |

**Table S11 – *A posteriori* contrasts for the effect of heatwave during male adulthood on female fertility for Dutch and Portuguese populations.** Reproductive success: number of adult offspring in each vial. *A posteriori* tukey contrasts. "T ratio": the T-test value obtained in each comparison. A) Comparison: Interaction between Treatment (Non-stressed + non-stressed, Stressed + non-stressed or Stressed + stressed) and Mating Opportunity (First or Second). B) Comparison: Interaction between Selection (Control or Warming) and Mating Opportunity (First or Second). Statistically significant terms are represented in bold.

A)

| History | Trait | Comparison | *T ratio* | *p-value* |
|---|---|---|---|---|
| Dutch | Reproductive Success | **First Non-stressed + non-stressed – First Stressed + non-stressed** | **5.038** | **< 0.001** |
| | | First Non-stressed + non-stressed – Second Stressed + non-stressed | -2.329 | 0.184 |
| | | **First Non-stressed + non-stressed – First Stressed + stressed** | **4.741** | **< 0.001** |
| | | First Non-stressed + non-stressed – Second Stressed + stressed | -1.648 | 0.567 |
| | | **First Non-stressed + non-stressed - Second Non-stressed + non-stressed** | **-5.689** | **< 0.001** |
| | | **Second Non-stressed + non-stressed – First Stressed + non-stressed** | **-9.194** | **< 0.001** |
| | | Second Non-stressed + non-stressed – Second Stressed + non-stressed | 2.546 | 0.112 |
| | | **Second Non-stressed + non-stressed – First Stressed + stressed** | **-8.804** | **< 0.001** |
| | | **Second Non-stressed + non-stressed - Second Stressed + stressed** | **3.114** | **0.023** |
| | | First Stressed + non-stressed – First Stressed + stressed | -0.217 | 1.000 |
| | | **First Stressed + non-stressed – Second Stressed + stressed** | **-6.478** | **< 0.001** |
| | | **First Stressed + non-stressed – Second Stressed + non-stressed** | **-8.012** | **< 0.001** |
| | | **Second Stressed + non-stressed – First Stressed + stressed** | **-6.789** | **< 0.001** |
| | | Second Stressed + non-stressed – Second Stressed + stressed | 0.649 | 0.987 |
| | | **First Stressed + stressed – Second Stressed + stressed** | **-6.901** | **< 0.001** |
| Portuguese | Reproductive Success | **First Non-stressed + non-stressed – First Stressed + non-stressed** | **4.983** | **< 0.001** |
| | | **First Non-stressed + non-stressed – Second Stressed + non-stressed** | **-2.835** | **0.009** |
| | | **First Non-stressed + non-stressed – First Stressed + stressed** | **6.561** | **< 0.001** |
| | | First Non-stressed + non-stressed – Second Stressed + stressed | -2.097 | 0.290 |
| | | **First Non-stressed + non-stressed - Second Non-stressed + non-stressed** | **-7.738** | **< 0.001** |
| | | **Second Non-stressed + non-stressed – First Stressed + non-stressed** | **-10.107** | **< 0.001** |
| | | **Second Non-stressed + non-stressed – Second Stressed + non-stressed** | **4.107** | **< 0.001** |
| | | **Second Non-stressed + non-stressed – First Stressed + stressed** | **-11.580** | **< 0.001** |
| | | **Second Non-stressed + non-stressed - Second Stressed + stressed** | **4.573** | **< 0.001** |
| | | First Stressed + non-stressed – First Stressed + stressed | 1.508 | 0.659 |
| | | **First Stressed + non-stressed – Second Stressed + stressed** | **-6.700** | **< 0.001** |
| | | **First Stressed + non-stressed – Second Stressed + non-stressed** | **-8.018** | **< 0.001** |
| | | **Second Stressed + non-stressed – First Stressed + stressed** | **-8.910** | **< 0.001** |
| | | Second Stressed + non-stressed – Second Stressed + stressed | 0.693 | 0.998 |
| | | **First Stressed + stressed – Second Stressed + stressed** | **-9.037** | **< 0.001** |

**B)**

| History | Trait | Comparison | T ratio | p-value |
|---|---|---|---|---|
| Portuguese | Reproductive Success | **First Control - First Warming** | **-3.798** | **< 0.001** |
| | | **First Control - Second Control** | **-11.663** | **< 0.001** |
| | | **First Control - Second Warming** | **-11.449** | **< 0.001** |
| | | **First Warming - Second Control** | **-6.814** | **< 0.001** |
| | | **First Warming - Second Warming** | **-8.889** | **< 0.001** |
| | | Second Control - Second Warming | -1.305 | 0.560 |

**Table S12 – Results from the analyses of variance of the effect of heatwave during male adulthood on female propensity to remate.** Remating: female mates with a different male after already mating once. Females with different bio-geographical origins ("History": Dutch or Portuguese), that were exposed for 45 generations to different thermal conditions ("Selection": Control or Warming), were paired twice with different males subjected, or not, to a heatwave treatment ("Treatment": Non-stressed + non-stressed, Stressed + non-stressed or Stressed + stressed) "Df": the degrees of freedom. "$X^2$": the Chi-square value obtained in each analysis. Statistically significant terms are represented in bold.

| Trait | Independent Variable | *Df* | $X^2$ | *p-value* |
|---|---|---|---|---|
| Remating | History | 1 | 3.362 | 0.067 |
| | Selection | 1 | 1.744 | 0.187 |
| | **Treatment** | **2** | **77.706** | **< 0.001** |

**Table S13 –*A posteriori* contrasts of the effect of heatwave during male adulthood on female propensity to remate.** Remating: female mates with a different male after already mating once. *A posteriori* tukey contrasts. "T ratio": the T-test value obtained in each comparison. A) Comparison between Treatments (Non-stressed + non-stressed, Stressed + non-stressed or Stressed + stressed). Statistically significant terms are represented in bold.

| Trait | Comparison | *T ratio* | *p-value* |
|---|---|---|---|
| Remating | **Non-stressed + non-stressed - Stressed + non-stressed** | **-8.736** | **< 0.001** |
| | **Non-stressed + non-stressed - Stressed + stressed** | **-7.191** | **< 0.001** |
| | **Stressed + non-stressed - Stressed + stressed** | **3.037** | **0.007** |

**Table S14 – Results from the analyses of variances of the effect of remating behaviour on the female reproductive success.** Reproductive success: number of adult offspring in each vial. Females paired twice with different males subjected, or not, to a heatwave treatment ("Treatment": Non-stressed + non-stressed, Stressed + non-stressed or Stressed + stressed), 1 and 3 days after the male thermal treatment ("Mating Opportunity": First or Second), displayed remating, or not ("Remating": No Remating or Remated). "Df": the degrees of freedom. "X2": the Chi-square value obtained in each analysis. Statistically significant terms are represented in bold.

| Trait | Treatment | Explanatory Variable | *Df* | $X^2$ | *p-value* |
|---|---|---|---|---|---|
| Reproductive Success | Non-stressed + non-stressed | **Mating Opportunity** | **1** | **100.527** | **< 0.001** |
| | | **Remating** | **1** | **4.808** | **0.028** |
| | Stressed + non-stressed | **Mating Opportunity** | **1** | **289.369** | **< 0.001** |
| | | **Remating** | **1** | **57.602** | **< 0.001** |
| | | **Mating Opportunity*Remating** | **1** | **40.931** | **< 0.001** |
| | Stressed + stressed | **Mating Opportunity** | **1** | **111.949** | **< 0.001** |
| | | **Remating** | **1** | **55.585** | **< 0.001** |
| | | **Mating Opportunity*Remating** | **1** | **45.504** | **< 0.001** |

**Table S15 – *A posteriori* contrasts for the effect of remating behaviour on the female reproductive success.** Reproductive success: number of adult offspring in each vial *A posteriori* tukey contrasts. “T ratio”: the T-test value obtained in each comparison. Comparison: Interaction between Remating (No Remating or Remated) and Mating Opportunity (First or Second). Statistically significant terms are represented in bold.

| Trait | Treatment | Comparison | *T ratio* | *p-value* |
|---|---|---|---|---|
| Reproductive Success | Stressed + non-stressed | **First No Remating - Second No Remating** | **-10.772** | **< 0.001** |
| | | **First No Remating - First Remated** | **7.514** | **< 0.001** |
| | | **First No Remating - Second Remated** | **-7.077** | **< 0.001** |
| | | **Second No Remating - First Remated** | **14.815** | **< 0.001** |
| | | **Second No Remating - Second Remated** | **3.426** | **0.004** |
| | | **First Remated - Second Remated** | **-14.139** | **< 0.001** |
| | Stressed + stressed | **First No Remating - Second No Remating** | **-7.100** | **< 0.001** |
| | | **First No Remating - First Remated** | **7.355** | **< 0.001** |
| | | **First No Remating - Second Remated** | **-3.999** | **0.004** |
| | | **Second No Remating - First Remated** | **9.478** | **< 0.001** |
| | | Second No Remating - Second Remated | 1.870 | 0.242 |
| | | **First Remated - Second Remated** | **-8.938** | **< 0.001** |

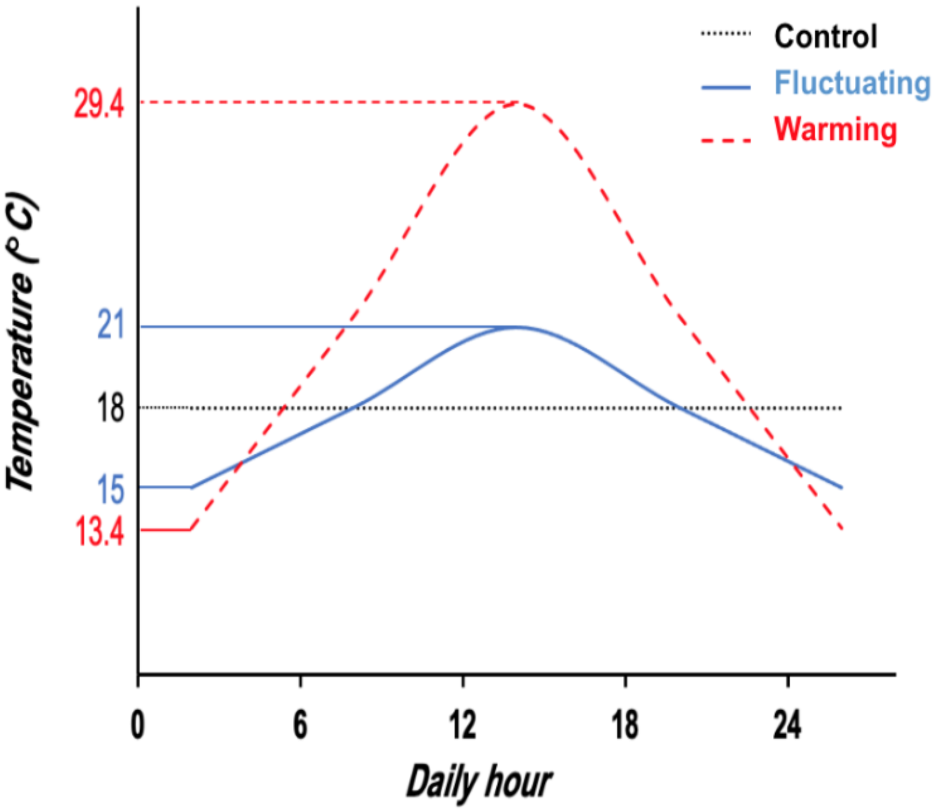


**Figure S1 – Daily temperature profile of the thermal regimes.** The black dashed line represents the Control regime, the red dashed line represents the Warming regime (from generation 20 of thermal evolution onwards), and the blue line the Fluctuating regime.

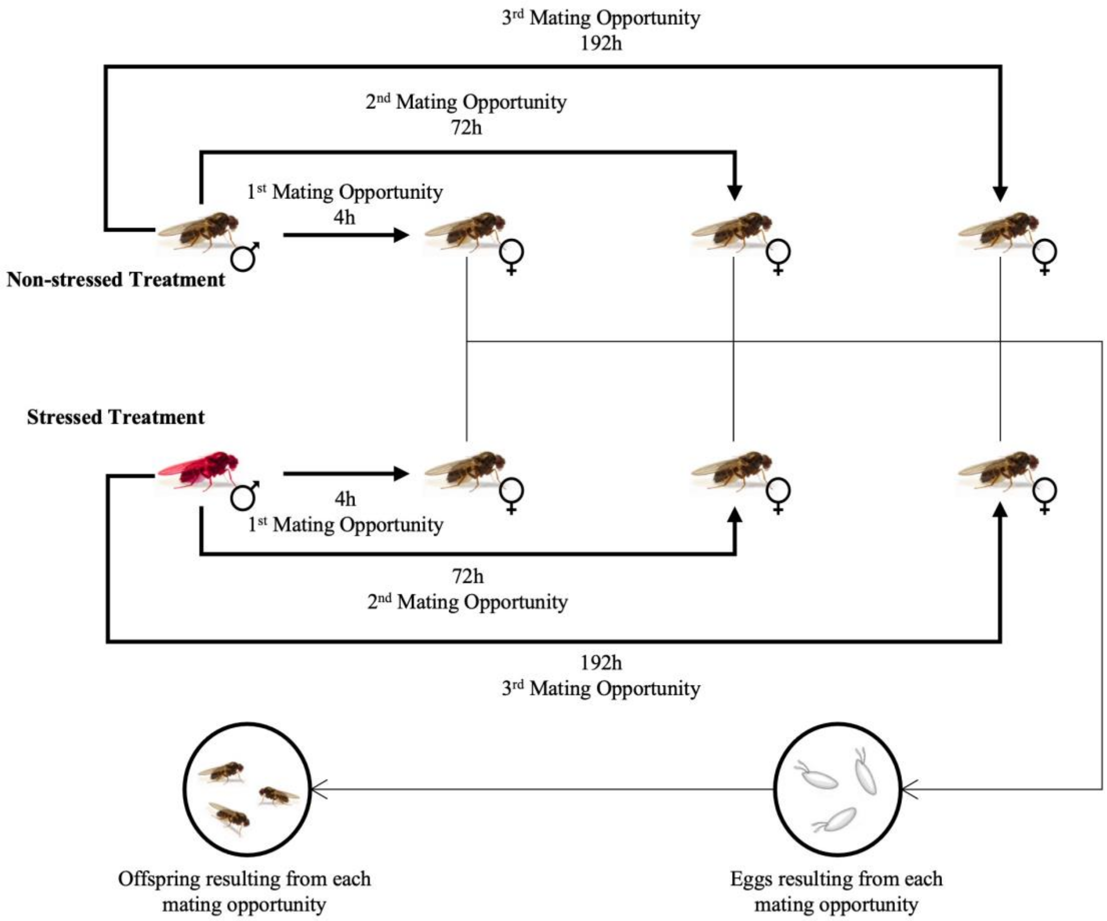


**Figure S2 - Schematic representation of the protocol used to test the recovery of male mating behaviour and fertility following heatwave.** The assay was performed using males from the Control and Warming regimes after 45 generations of thermal evolution. Red flies represent stressed males (exposed to a heatwave of 31°C for 69 hours), while non-coloured flies represent non-stressed males and females (adapted from Mestres et al., 2016). The time points represent the hours after heatwave when each mating opportunity occurred. In each mating opportunity the courtship latency, copulation latency, and copulation duration were registered for 2 hours. Later, for each vial, the reproductive success was assessed by measuring the number of adult offspring produced.

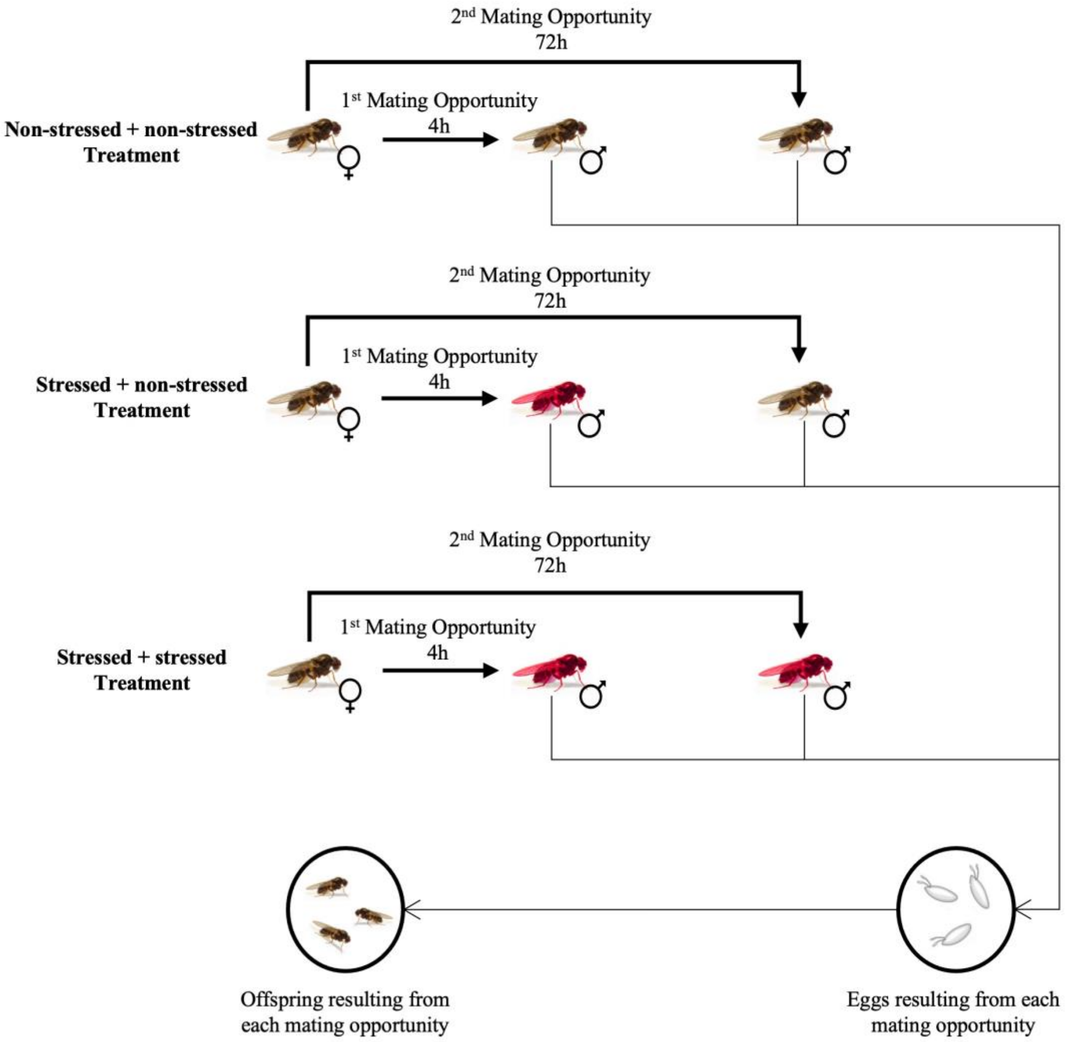


**Figure S3 - Schematic representation of the protocol used to test the female mating behaviour and fertility following male heatwave.** The assay was performed in females from the Control and Warming regimes after 45 generations of thermal evolution. Red flies represent stressed males (exposed to a heatwave of 31°C for 69 hours), while non-coloured flies represent non-stressed males and females (adapted from Mestres et al., 2016). The time points represent the hours after heatwave when each mating opportunity occurred. In each mating opportunity the courtship latency, copulation latency, and copulation duration were registered for 2 hours. Later, for each vial, the reproductive success was assessed by measuring the number of adult offspring produced.
