## Supplement 1 for "Monandrous flies do remate: plastic and evolutionary consequences of heat exposure on mating behaviour and fertility in *Drosophila subobscura*"

2 - Universidade de Lisboa, Lisbon, Portugal. Departamento de Biologia Animal, Faculdade de Ciências.

3 - Egas Moniz Center for Interdisciplinary Research (CiiEM), Egas Moniz School of Health and Science, Caparica, Portugal.

* - Co-last authors.

### Supplement 1

#### **Material and methods**

##### ***Statistical analyses***

###### *Recovery of male fertility following heatwave*

To analyse how a heatwave during early life influenced males' fertility, two traits were studied: fecundity and offspring viability.

To assess fecundity (number of eggs per vial) a GLMM model with a quasi-Poisson error distribution and a parameter to account for zero-inflated data (ziformula ~1; package *glmmTMB*) was used. Offspring viability was computed using the function *cbind* with the number of successfully hatched eggs and the number of unhatched eggs as arguments and was analysed using a GLMM model with a binomial error distribution. The models for these two traits included History, Selection, Treatment, Mating Opportunity, and the interaction between them as fixed factors. Block, defined as the set of same numbered replicate populations (1, 2, or 3), was included as a random factor. Given that each male was tested multiple times (throughout the three mating opportunities), we accounted for repeated measures (Park et al., 2009) by adding the interaction between male ID (unique identification of each male) and Mating Opportunity as a covariate to the model. Due to a significant quadruple interaction between all factors affecting these traits, analyses were done separately for populations from different bio-geographical origins (factor History). The analyses were similar to that described but without History as fixed factor. The error structure of these models was also similar to that of the original models, except for the analysis of fecundity of Dutch populations, where the model used was a GLMM with a Poisson error distribution and a parameter to account for zero inflation (ziformula ~1). Furthermore, these models had a significant triple interaction with the Treatment factor was obtained, so additional analyses were done separating the different treatments (Non-stressed or Stressed). These models included Selection, Mating Opportunity, and their interaction as fixed factors and Block as a random factor. The models for the analysis of fecundity were GLMM models with a Poisson error distribution and a parameter to account for zero inflation (ziformula ~1), while for offspring viability the models were GLMM models with a binomial error distribution.

*Female fertility following male heatwave*

To analyse how female fertility was affected by male thermal treatment (stressed or non-stressed), two traits were studied: fecundity and offspring viability.

Fecundity was analysed using a GLMM model with a quasi-Poisson error distribution and a parameter to account for zero inflation (ziformula ~1). Offspring viability was computed as previously described and was analysed using a GLMM model with a binomial error distribution. The models for these traits included History, Selection, Treatment, Mating Opportunity, and the interaction between them as fixed factors. Block, defined as the set of same numbered replicate populations (1, 2, or 3), was included as a random factor. Given that the same females were tested in two different moments (first or second mating opportunities), we accounted for repeated measures by adding the interaction between female ID (unique identification of each female) and Mating Opportunity as covariate to the model. Due to multiple significant interactions involving the factor History in the analyses of fecundity, a new analysis for this trait was done separately for populations with distinct bio-geographical origins. The model was identical to that described above but without History as fixed factor. The error structure of the model was also similar, except for the analysis of fecundity of Dutch populations, where the model used was a GLMM with a Poisson error distribution and a parameter to account for zero inflation (ziformula ~1).

#### Results

##### *Effect of heatwave on male fecundity and offspring viability*

The male fecundity of Dutch populations was significantly affected by a triple interaction between Selection, Treatment and Mating Opportunity ($X^2$ = 7.677, p = 0.022; Table 1 Supplement 1). Analyses done for each treatment separately showed that non-stressed Dutch males from both selective regimes had a similar response throughout mating opportunities, with Mating opportunity being the only significant factor ($X^2$ = 13.010, p = 0.001; see Table 2 Supplement 1), corresponding to a significant increase in fecundity from the second to the third mating opportunity (see Table 3A Supplement, Figure 1A Supplement 1). On the other hand, stressed males from Dutch populations displayed a distinct dynamic between mating opportunities depending on their selection regime (Selection x Mating Opportunity, $X^2 = 6.346$, $p = 0.042$; see Table 2 Supplement 1, Figure 1A Supplement 1). Despite this significant interaction, *a posteriori* contrasts did not show any differences in fecundity across mating opportunities between populations from different selective regimes (see Table 3 Supplement 1, Figure 1A Supplement 1). The fecundity of populations from Portugal was significantly affected by the interaction between Treatment and Mating Opportunity ($X^2 = 7.102$, $p = 0.029$; see Table 1 Supplement 1), with males from different treatments showing distinct dynamics across mating opportunities. Indeed, comparing the first and the last (third) mating opportunities, a slight increase in fecundity was observed for stressed males but not for the non-stressed ones (see Table 4 Supplement 1, Figure 1A Supplement 1). However, differences between males from distinct treatments remained significant in the last mating opportunity (see Table 4 Supplement 1, Figure 1A Supplement 1), with non-stressed males displaying higher levels of fecundity.

The offspring viability of both Dutch and Portuguese populations was affected by a significant interaction between Selection, Treatment and Mating Opportunity ($X^2 = 7.808$, $p = 0.020$, for Dutch populations; $X^2 = 6.067$, $p = 0.048$, for Portuguese populations; see Table 1 Supplement 1). Given this triple interaction, additional analyses were performed for each Treatment separately for both the Dutch and Portuguese populations. The offspring viability of both non-stressed and stressed Dutch males, was significantly shaped by the Mating Opportunity ($X^2 = 6.112$, $p = 0.047$, for non-stressed males; $X^2 = 22.222$, $p < 0.001$, for stressed males; see Table 2 Supplement 1), with non-stressed and stressed males presenting a similar pattern. The offspring viability of both types of males was lower in the third relative to the second mating opportunity (see Table 3A Supplement 1, Figure 1B Supplement 1). For the offspring viability of Portuguese non-stressed males, the Mating Opportunity was once again the only significant factor ($X^2$ = 8.470, p = 0.014; see Table 2 Supplement 1), with a decrease in offspring viability being observed in the last mating opportunity relative to the first one (see Table 3A Supplement 1, Figure 1B Supplement 1). In turn, the offspring viability of Portuguese stressed males was influenced by an interaction between Selection and Mating Opportunity ($X^2$ = 8.894, p = 0.012; Table 2 Supplement 1): Indeed, stressed males from the Warming regime had a significant increase in offspring viability from the first to the second mating opportunity (see Table 3B Supplement 1), and kept similar levels of offspring viability in the third mating opportunity when compared to the second one. On the other hand, stressed Portuguese males from the Control regime, had similar offspring viability in the first and second mating opportunities, but a decrease in the final mating opportunity (see Table 3B Supplement 1, Figure 1B Supplement 1).

*Effect of reduced male fertility on female fecundity and offspring viability*

Multiple interactions with the variable History had a significant effect on fecundity (see Table 5 Supplement 1), thus statistical analyses for Dutch and Portuguese populations were performed separately for this trait. The fecundity of both Dutch and Portuguese populations was significantly shaped by the interaction between Treatment and Mating Opportunity ($X^2$ = 14.311, p < 0.001; $X^2$ = 10.814, p = 0.004, respectively; see Table 6 Supplement 1). In the first mating opportunity, females that first mated with males subjected to a heatwave had a significant reduction in their fecundity relative to those that mated with non-stressed males (see Table 7A Supplement 1, Figure 2A Supplement 1). While an increase in fecundity was observed between mating opportunities for all treatments, this increase was significantly steeper in females that first had the chance to mate with stressed males compared to females paired with two non-stressed males. In Dutch populations, the fecundity of all females was similar in the second mating opportunity, revealing that Dutch females that first had the opportunity to mate with a male exposed to high temperatures were able to fully rescue their fecundity after a second mating opportunity, regardless of the thermal treatment of the second male (see Table 7A Supplement 1, Figure 2A Supplement 1). However, in Portuguese populations, this increase in fecundity from the first to the second mating, was not enough to reach the fecundity levels of females that were paired with two non-stressed males (see Table 7A Supplement 1, Figure 2A Supplement 1). Additionally, the fecundity of Portuguese populations was also affected by an interaction between Selection and Mating Opportunity ($X^2$ = 9.197, p = 0.002; see Table 6 Supplement 1), with females from the Control regime having lower fecundity in the first mating opportunity than females from the Warming regime, but similar levels in the second mating opportunity (see Table 7B Supplement 1, Figure 2A Supplement 1).

The offspring viability was significantly shaped by the interaction between Treatment and Mating Opportunity interaction ($X^2$ = 18.208, p < 0.001; see Table 8 Supplement 1), but not by History or Selection, with different treatments presenting distinct dynamics across mating opportunities. Indeed, offspring viability was significantly lower in the first mating opportunity when females mated with stressed males relative to females that first mated with a non-stressed male (see Table 9 Supplement 1, Figure 2B Supplement 1). However, this was not true for the second mating opportunity, with females that had the chance to mate with two non-stressed males having similar offspring viability in both mating opportunities (see Table 9 Supplement 1), while females that first mated with stressed males significantly increased their offspring viability from the first to the second mating opportunity, regardless of the thermal treatment of the second male. In fact, the offspring viability after the second mating opportunity was similar across all treatments (see Table 9 Supplement). This pattern shows that a second mating opportunity for females that first mated with a stressed male can rescue their offspring viability (see Figure 2B).

#### Tables

**Table 1 Supplement 1 – Results from the analyses of variances of the effect of heatwave during male adulthood on male fertility when analysing populations of different history separately.** Fecundity: Number of laid eggs; Offspring viability: Ratio between the number of offspring (adult offspring) and the number of laid eggs (total number of eggs). Males with different bio-geographical origins ("History": Dutch or Portuguese) that were exposed for 45 generations to different thermal conditions ("Selection": Control or Warming), were subjected, or not, to a heatwave treatment ("Treatment": Non-stressed or Stressed) and were paired, with different females, 1, 3 and 8 days after the thermal treatment ("Mating Opportunity": First, Second or Third). "Df": the degrees of freedom. "$X^2$": the Chi-square value obtained in each analysis. Statistically significant terms are represented in bold.

| History | Trait | Independent Variable | *Df* | $X^2$ | *p-value* |
|---|---|---|---|---|---|
| Dutch | Fecundity | Selection | 1 | 0.299 | 0.585 |
| | | **Treatment** | **1** | **40.770** | **< 0.001** |
| | | **Mating Opportunity** | **2** | **8.306** | **0.016** |
| | | Selection*Treatment | 1 | 0.792 | 0.374 |
| | | Selection*Mating Opportunity | 2 | 4.098 | 0.129 |
| | | Treatment*Mating Opportunity | 2 | 2.564 | 0.277 |
| | | **Selection*Treatment*Mating Opportunity** | **2** | **7.677** | **0.022** |
| | Offspring viability | Selection | 1 | 0.452 | 0.501 |
| | | **Treatment** | **1** | **52.626** | **< 0.001** |
| | | **Mating Opportunity** | **2** | **28.135** | **< 0.001** |
| | | Selection*Treatment | 1 | 1.142 | 0.285 |
| | | Selection*Mating Opportunity | 2 | 0.105 | 0.949 |
| | | **Treatment*Mating Opportunity** | **2** | **8.252** | **0.016** |
| | | **Selection*Treatment*Mating Opportunity** | **2** | **7.808** | **0.020** |
| Portuguese | Fecundity | Selection | 1 | 1.069 | 0.301 |
| | | **Treatment** | **1** | **36.120** | **< 0.001** |
| | | **Mating Opportunity** | **2** | **13.794** | **0.001** |
| | | **Treatment*Mating Opportunity** | **2** | **7.102** | **0.029** |
| | Offspring viability | Selection | 1 | 1.158 | 0.282 |
| | | **Treatment** | **1** | **124.425** | **< 0.001** |
| | | **Mating Opportunity** | **2** | **21.730** | **< 0.001** |
| | | **Selection*Treatment** | **1** | **4.442** | **0.035** |
| | | **Selection*Mating Opportunity** | **2** | **13.502** | **0.001** |
| | | **Treatment*Mating Opportunity** | **2** | **24.379** | **< 0.001** |
| | | **Selection*Treatment*Mating Opportunity** | **2** | **6.067** | **0.048** |

**Table 2 Supplement 1 –Results from the analysis of variance of the effect of heatwave during male adulthood on male fertility for Dutch and Portuguese populations separated by treatment (Non-stressed or Stressed).** Fecundity: Number of laid eggs; Offspring viability: Ratio between the number of offspring (adult offspring) and the number of laid eggs (total number of eggs). Males with different bio-geographical origins ("History": Dutch or Portuguese) that were exposed for 45 generations to different thermal conditions ("Selection": Control or Warming), were subjected, or not, to a heatwave treatment ("Treatment": Non-stressed or Stressed) and were paired, with different females, 1, 3 and 8 days after the thermal treatment ("Mating Opportunity": First, Second or Third). "Df": the degrees of freedom. "$X^2$": the Chi-square value obtained in each analysis. Statistically significant terms are represented in bold.

| History | Treatment | Trait | Explanatory Variable | *Df* | $X^2$ | *p-value* |
|---|---|---|---|---|---|---|
| Dutch | Non-Stressed | Fecundity | Selection | 1 | 0.045 | 0.832 |
| | | | **Mating Opportunity** | **2** | **13.010** | **0.001** |
| | | Offspring viability | Selection | 1 | 0.033 | 0.857 |
| | | | **Mating Opportunity** | **2** | **6.112** | **0.047** |
| | | | Selection*Mating Opportunity | 2 | 5.132 | 0.077 |
| | Stressed | Fecundity | Selection | 1 | 0.456 | 0.499 |
| | | | Mating Opportunity | 2 | 0.663 | 0.718 |
| | | | **Selection*Mating Opportunity** | **2** | **6.346** | **0.042** |
| | | Offspring viability | Selection | 1 | 0.220 | 0.640 |
| | | | **Mating Opportunity** | **2** | **22.222** | **< 0.001** |
| Portuguese | Non-Stressed | Offspring viability | Selection | 1 | 3.757 | 0.053 |
| | | | **Mating Opportunity** | **2** | **8.470** | **0.014** |
| | Stressed | Offspring viability | Selection | 1 | 2.589 | 0.108 |
| | | | **Mating Opportunity** | **2** | **21.317** | **< 0.001** |
| | | | **Selection*Mating Opportunity** | **2** | **8.894** | **0.012** |

**Table 3 Supplement 1 –*A posteriori* contrasts of the effect of heatwave during male adulthood on male fecundity and offspring viability for Dutch and Portuguese populations separated by treatment (Non-stressed or Stressed).** Fecundity: Number of laid eggs; Offspring viability: Ratio between the number of offspring (adult offspring) and the number of laid eggs (total number of eggs). *A posteriori* tukey contrasts. "T ratio": the T-test value obtained in each comparison. A) Comparison between Mating Opportunities (First, Second or Third). B) Comparison: Interaction between Selection (Control or Warming) and Mating Opportunity (First, Second or Third). Statistically significant terms are represented in bold.

A)

| History | Treatment | Trait | Comparison | *T ratio* | *p-value* |
|---|---|---|---|---|---|
| Dutch | Non-Stressed | Fecundity | First - Second | 2.181 | 0.076 |
| | | | First - Third | -2.028 | 0.107 |
| | | | **Second - Third** | **-3.603** | **0.001** |
| | | Offspring viability | First - Second | -1.820 | 0.165 |
| | | | First - Third | 1.034 | 0.556 |
| | | | **Second - Third** | **2.450** | **0.039** |
| | Stressed | Offspring viability | First - Second | -1.643 | 0.229 |
| | | | First - Third | 2.297 | 0.057 |
| | | | **Second - Third** | **4.711** | **< 0.001** |
| Portuguese | Non-stressed | Offspring viability | First - Second | 1.158 | 0.479 |
| | | | **First - Third** | **2.863** | **0.012** |
| | | | Second - Third | 1.960 | 0.124 |

B)

| History | Treatment | Trait | Comparison | *T ratio* | *p-value* |
|---|---|---|---|---|---|
| Dutch | Stressed | Fecundity | Control x First - Warming x First | -0.594 | 0.991 |
| | | | Control x First - Warming x Second | -0.448 | 0.998 |
| | | | Control x First - Warming x Third | 0.432 | 0.998 |
| | | | Control x First - Control x Second | -0.107 | 1.000 |
| | | | Control x First - Control x Third | -1.919 | 0.392 |
| | | | Control x Second - Warming x First | 0.501 | 0.996 |
| | | | Control x Second - Warming x Second | -0.355 | 0.999 |
| | | | Control x Second - Warming x Third | 0.528 | 0.995 |
| | | | Control x Second - Control x Third | -2.305 | 0.1947 |
| | | | Control x Third - Warming x First | -1.243 | 0.816 |
| | | | Control x Third - Warming x Second | -1.465 | 0.687 |
| | | | Control x Third - Warming x Third | 2.425 | 0.150 |
| | | | Warming x First - Warming x Second | 0.174 | 1.000 |
| | | | Warming x First - Warming x Third | 0.970 | 0.927 |
| | | | Warming x Second - Warming x Third | 1.041 | 0.904 |
| Portuguese | Stressed | Offspring viability | Control x First - Warming x First | 2.370 | 0.169 |
| | | | Control x First - Warming x Second | -0.250 | 1.000 |
| | | | Control x First - Warming x Third | 0.488 | 0.997 |
| | | | Control x First - Control x Second | -2.170 | 0.254 |
| | | | Control x First - Control x Third | 1.692 | 0.538 |
| | | | **Control x Second - Warming x First** | **-4.455** | **< 0.001** |
| | | | Control x Second - Warming x Second | 2.124 | 0.277 |
| | | | Control x Second - Warming x Third | 2.638 | 0.090 |
| | | | **Control x Second - Control x Third** | **4.222** | **< 0.001** |
| | | | Control x Third - Warming x First | -0.976 | 0.925 |
| | | | Control x Third - Warming x Second | 2.087 | 0.296 |
| | | | Control x Third - Warming x Third | -1.114 | 0.876 |
| | | | **Warming x First - Warming x Second** | **-3.262** | **0.015** |
| | | | Warming x First - Warming x Third | -2.204 | 0.238 |
| | | | Warming x Second - Warming x Third | 0.946 | 0.934 |

**Table 4 Supplement 1 – *A posteriori* contrasts for the effect of heatwave during male adulthood on male fecundity for Portuguese populations.** Fecundity: Number of laid eggs. *A posteriori* tukey contrasts. “T ratio”: the T-test value obtained in each comparison. Comparison: Interaction between Treatment (Stressed or Non-stressed) and Mating Opportunity (First, Second or Third). Statistically significant terms are represented in bold.

| History | Trait | Comparison | *T ratio* | *p-value* |
|---|---|---|---|---|
| Portuguese | Fecundity | **Non-stressed x First - Stressed x First** | **5.050** | **<0.001** |
| | | Non-stressed x First - Stressed x Second | 2.804 | 0.058 |
| | | Non-stressed x First - Stressed x Third | 2.345 | 0.177 |
| | | Non-stressed x First - Non-stressed x Second | 1.529 | 0.646 |
| | | Non-stressed x First - Non-stressed x Third | -2.023 | 0.330 |
| | | **Non-stressed x Second - Stressed x First** | **-3.813** | **0.002** |
| | | Non-stressed x Second - Stressed x Second | 1.524 | 0.649 |

| | | |
|---|---|---|
| Non-stressed x Second - Stressed x Third | 0.966 | 0.929 |
| **Non-stressed x Second - Non-stressed x Third** | **-3.494** | **0.007** |
| **Non-stressed x Third - Stressed x First** | **-6.812** | **<0.001** |
| **Non-stressed x Third - Stressed x Second** | **-4.575** | **0.001** |
| **Non-stressed x Third - Stressed x Third** | **4.247** | **0.003** |
| Stressed x First - Stressed x Second | -2.158 | 0.259 |
| **Stressed x First - Stressed x Third** | **-2.889** | **0.046** |
| Stressed x Second - Stressed x Third | -0.582 | 0.992 |

**Table 5 Supplement 1 – Results from the analyses of variances of the effect of heatwave during male adulthood on female fecundity.** Fecundity: Number of laid eggs. Females with different bio-geographical origins ("History": Dutch or Portuguese), that were exposed for 45 generations to different thermal conditions ("Selection": Control or Warming), were paired twice with different males subjected, or not, to a heatwave treatment ("Treatment": Non-stressed + non-stressed, Stressed + non-stressed or Stressed + stressed), 1 and 3 days after the male thermal treatment ("Mating Opportunity": First or Second). "Df": the degrees of freedom. Df.res: residual degrees of freedom. "$X^2$": the Chi-square value obtained in each analysis. Statistically significant terms are represented in bold.

| Trait | Explanatory Variable | *Df* | $X^2$ | *p-value* |
|---|---|---|---|---|
| Fecundity | History | 1 | 0.814 | 0.367 |
| | **Selection** | **1** | **6.304** | **0.012** |
| | **Treatment** | **2** | **87.430** | **< 0.001** |
| | **Mating Opportunity** | **1** | **357.253** | **< 0.001** |
| | **History*Selection** | **1** | **5.497** | **0.019** |
| | **History*Mating Opportunity** | **1** | **10.035** | **0.002** |
| | **Selection*Mating Opportunity** | **1** | **10.559** | **0.001** |
| | **Treatment*Mating Opportunity** | **2** | **20.950** | **< 0.001** |

**Table 6 Supplement 1 – Results from the analyses of variances of the effect of heatwave during male adulthood on female fecundity when analysing populations of different history separately.** Fecundity: Number of laid eggs. Females with different bio-geographical origins ("History": Dutch or Portuguese), that were exposed for 45 generations to different thermal conditions ("Selection": Control or Warming), were paired twice with different males subjected, or not, to a heatwave treatment ("Treatment": Non-stressed + non-stressed, Stressed + non-stressed or Stressed + stressed), 1 and 3 days after the male thermal treatment ("Mating Opportunity": First or Second). "Df": the degrees of freedom. Df.res: residual degrees of freedom. "$X^2$": the Chi-square value obtained in each analysis. Statistically significant terms are represented in bold.

| History | Trait | Explanatory Variable | *Df* | $X^2$ | *p-value* |
|---|---|---|---|---|---|
| Dutch | Fecundity | Selection | 1 | 0.465 | 0.495 |
| | | **Treatment** | **2** | **33.490** | **< 0.001** |
| | | **Mating Opportunity** | **1** | **143.136** | **< 0.001** |
| | | **Treatment*Mating Opportunity** | **2** | **14.311** | **< 0.001** |
| Portuguese | Fecundity | **Selection** | **1** | **12.476** | **< 0.001** |
| | | **Treatment** | **2** | **54.232** | **< 0.001** |
| | | **Mating Opportunity** | **1** | **204.534** | **< 0.001** |
| | | **Selection*Mating Opportunity** | **1** | **9.197** | **0.002** |
| | | **Treatment*Mating Opportunity** | **2** | **10.814** | **0.004** |

**Table 7 Supplement 1 – *A posteriori* contrasts for the effect of heatwave during male adulthood on female fecundity for Dutch and Portuguese populations.** Fecundity: Number of laid eggs. *A posteriori* tukey contrasts. "T ratio": the T-test value obtained in each comparison. A) Comparison: Interaction between Treatment (Non-stressed x non-stressed, Stressed x non-stressed or Stressed x stressed) and Mating Opportunity (First or Second). B) Comparison:

Interaction between Selection (Control or Warming) and Mating Opportunity (First or Second). Statistically significant terms are represented in bold.

A)

| History | Trait | Comparison | *T ratio* | *p-value* |
|---|---|---|---|---|
| Dutch | Fecundity | **Non-stressed x non-stressed x First – Stressed x non-stressed x First** | **4.989** | **< 0.001** |
| | | Non-stressed x non-stressed x First – Stressed x non-stressed x Second | -1.738 | 0.507 |
| | | **Non-stressed x non-stressed x First – Stressed x stressed x First** | **4.817** | **< 0.001** |
| | | Non-stressed x non-stressed x First – Stressed x stressed x Second | -1.424 | 0.713 |
| | | **Non-stressed x non-stressed x First - Non-stressed x non-stressed x Second** | **-4.626** | **< 0.001** |
| | | **Non-stressed x non-stressed x Second – Stressed x non-stressed x First** | **-9.000** | **< 0.001** |
| | | Non-stressed x non-stressed x Second – Stressed x non-stressed x Second | 2.548 | 0.112 |
| | | **Non-stressed x non-stressed x Second – Stressed x stressed x First** | **-8.703** | **< 0.001** |
| | | Non-stressed x non-stressed x Second - Stressed x stressed x Second | 2.799 | 0.059 |
| | | Stressed x non-stressed x First – Stressed x stressed x First | -0.079 | 1.000 |
| | | **Stressed x non-stressed x First – Stressed x stressed x Second** | **-6.772** | **< 0.001** |
| | | **Stressed x non-stressed x First – Stressed x non-stressed x Second** | **-8.268** | **< 0.001** |
| | | **Stressed x non-stressed x Second – Stressed x stressed x First** | **-6.891** | **< 0.001** |
| | | Stressed x non-stressed x Second – Stressed x stressed x Second | 0.335 | 0.999 |
| | | **Stressed x stressed x First – Stressed x stressed x Second** | **-7.616** | **< 0.001** |
| Portuguese | Fecundity | **Non-stressed x non-stressed x First – Stressed x non-stressed x First** | **4.914** | **< 0.001** |
| | | **Non-stressed x non-stressed x First – Stressed x non-stressed x Second** | **-3.417** | **0.009** |
| | | **Non-stressed x non-stressed x First – Stressed x stressed x First** | **5.543** | **< 0.001** |
| | | **Non-stressed x non-stressed x First – Stressed x stressed x Second** | **-2.935** | **0.040** |
| | | **Non-stressed x non-stressed x First - Non-stressed x non-stressed x Second** | **-8.016** | **< 0.001** |
| | | **Non-stressed x non-stressed x Second – Stressed x non-stressed x First** | **-10.736** | **< 0.001** |
| | | **Non-stressed x non-stressed x Second – Stressed x non-stressed x Second** | **4.205** | **< 0.001** |
| | | **Non-stressed x non-stressed x Second – Stressed x stressed x First** | **-11.201** | **< 0.001** |
| | | **Non-stressed x non-stressed x Second - Stressed x stressed x Second** | **4.383** | **< 0.001** |
| | | Stressed x non-stressed x First – Stressed x stressed x First | 0.674 | 0.985 |
| | | **Stressed x non-stressed x First – Stressed x stressed x Second** | **-7.535** | **< 0.001** |
| | | **Stressed x non-stressed x First – Stressed x non-stressed x Second** | **-8.685** | **< 0.001** |
| | | **Stressed x non-stressed x Second – Stressed x stressed x First** | **-8.559** | **< 0.001** |
| | | Stressed x non-stressed x Second – Stressed x stressed x Second | 0.420 | 0.998 |
| | | **Stressed x stressed x First – Stressed x stressed x Second** | **-8.817** | **< 0.001** |

B)

| History | Trait | Comparison | T ratio | p-value |
|---|---|---|---|---|

| | | | | |
|---|---|---|---|---|
| Portuguese | Fecundity | **Control x First - Warming x First** | **-3.811** | **< 0.001** |
| | | **Control x First - Control x Second** | **-12.125** | **< 0.001** |
| | | **Control x First - Warming x Second** | **-11.838** | **< 0.001** |
| | | **Warming x First - Control Second** | **-7.094** | **< 0.001** |
| | | **Warming x First - Warming x Second** | **-8.925** | **< 0.001** |
| | | Control x Second - Warming x Second | -1.164 | 0.650 |

**Table 8 Supplement 1– Results from the analyses of variances of the effect of heatwave during male adulthood on female offspring viability.** Offspring viability: Ratio between the number of offspring (adult offspring) and the number of laid eggs (total number of eggs). Females with different bio-geographical origins (“History”: Dutch or Portuguese), that were exposed for 45 generations to different thermal conditions (“Selection”: Control or Warming), were paired twice with different males subjected, or not, to a heatwave treatment (“Treatment”: Non-stressed + non-stressed, Stressed + non-stressed or Stressed + stressed), 1 and 3 days after the male thermal treatment (“Mating Opportunity”: First or Second). “Df”: the degrees of freedom. Df.res: residual degrees of freedom. “$X^2$”: the Chi-square value obtained in each analysis. Statistically significant terms are represented in bold.

| Trait | Independent Variable | *Df* | $X^2$ | *p-value* |
|---|---|---|---|---|
| Offspring viability | History | 1 | 3.400 | 0.0652 |
| | Selection | 1 | 0.075 | 0.7836 |
| | **Treatment** | **2** | **42.661** | **< 0.001** |
| | **Mating Opportunity** | **1** | **30.537** | **< 0.001** |
| | **Treatment*Mating Opportunity** | **2** | **18.208** | **< 0.001** |

**Table 9 Supplement – *A posteriori* contrasts for the effect of heatwave during male adulthood on female offspring viability.** Offspring viability: Ratio between the number of offspring (adult offspring) and the number of laid eggs (total number of eggs). *A posteriori* tukey contrasts. “T ratio”: the T-test value obtained in each comparison. Comparison: Interaction between Treatment (Non-stressed x non-stressed, Stressed x non-stressed or Stressed x stressed) and Mating Opportunity (First or Second). Statistically significant terms are represented in bold.

| Trait | Comparison | *T ratio* | *p-value* |
|---|---|---|---|
| Offspring viability | **Non-stressed x non-stressed x First – Stressed x non-stressed x First** | **6.531** | **< 0.001** |
| | Non-stressed x non-stressed x First – Stressed x non-stressed x Second | 1.816 | 0.455 |
| | **Non-stressed x non-stressed x First – Stressed x stressed x First** | **6.157** | **< 0.001** |
| | Non-stressed x non-stressed x First – Stressed x stressed x Second | 2.546 | 0.112 |
| | Non-stressed x non-stressed x First - Non-stressed x non-stressed x Second | -0.008 | 1.000 |
| | **Non-stressed x non-stressed x Second – Stressed x non-stressed x First** | **-6.445** | **< 0.001** |
| | Non-stressed x non-stressed x Second – Stressed x non-stressed x Second | 1.821 | 0.452 |
| | **Non-stressed x non-stressed x Second – Stressed x stressed x First** | **-6.076** | **< 0.001** |
| | Non-stressed x non-stressed x Second - Stressed x stressed x Second | 2.550 | 0.111 |
| | Stressed x non-stressed x First – Stressed x stressed x First | -0.301 | 1.00 |
| | **Stressed x non-stressed x First – Stressed x stressed x Second** | **-3.971** | **0.001** |
| | **Stressed x non-stressed x First – Stressed x non-stressed x Second** | **-5.403** | **< 0.001** |
| | **Stressed x non-stressed x Second – Stressed x stressed x First** | **-4.471** | **0.001** |
| | Stressed x non-stressed x Second – Stressed x stressed x Second | 0.796 | 0.968 |
| | **Stressed x stressed x First – Stressed x stressed x Second** | **-4.108** | **< 0.001** |

### Figures

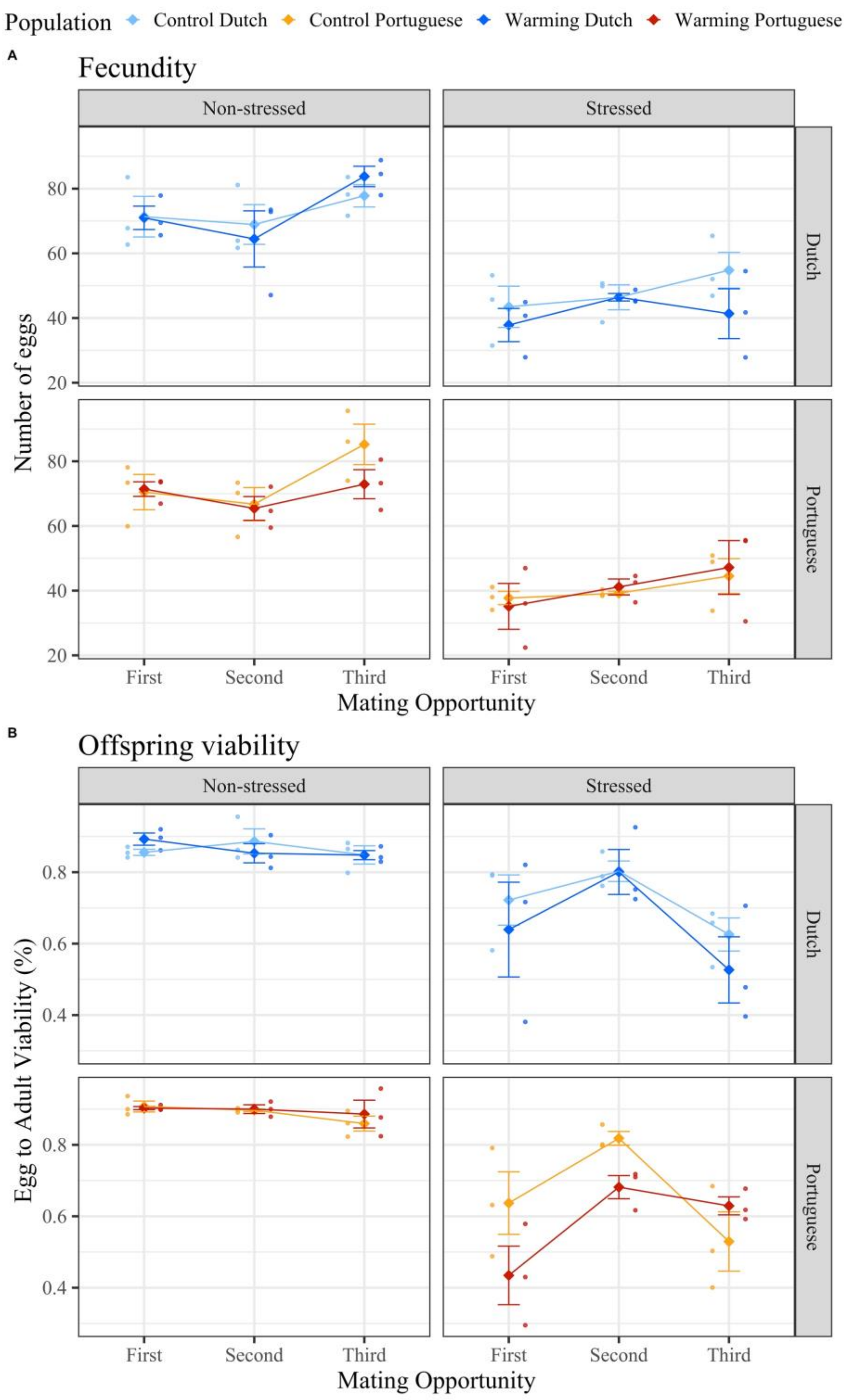


**Figure 1 – Effect of heatwave during male adulthood on male fertility.** A) Fecundity: number of laid eggs; B) Offspring viability: Ratio between the number of offspring and the number of laid eggs. Colder colours represent Dutch populations, while warmer ones represent Portuguese populations. Lighter tones represent the Control regime, while darker tones represent the Warming regime. The small circles represent the mean values of each replicate population, and the big diamonds represent the mean of the three replicate populations (±SE).

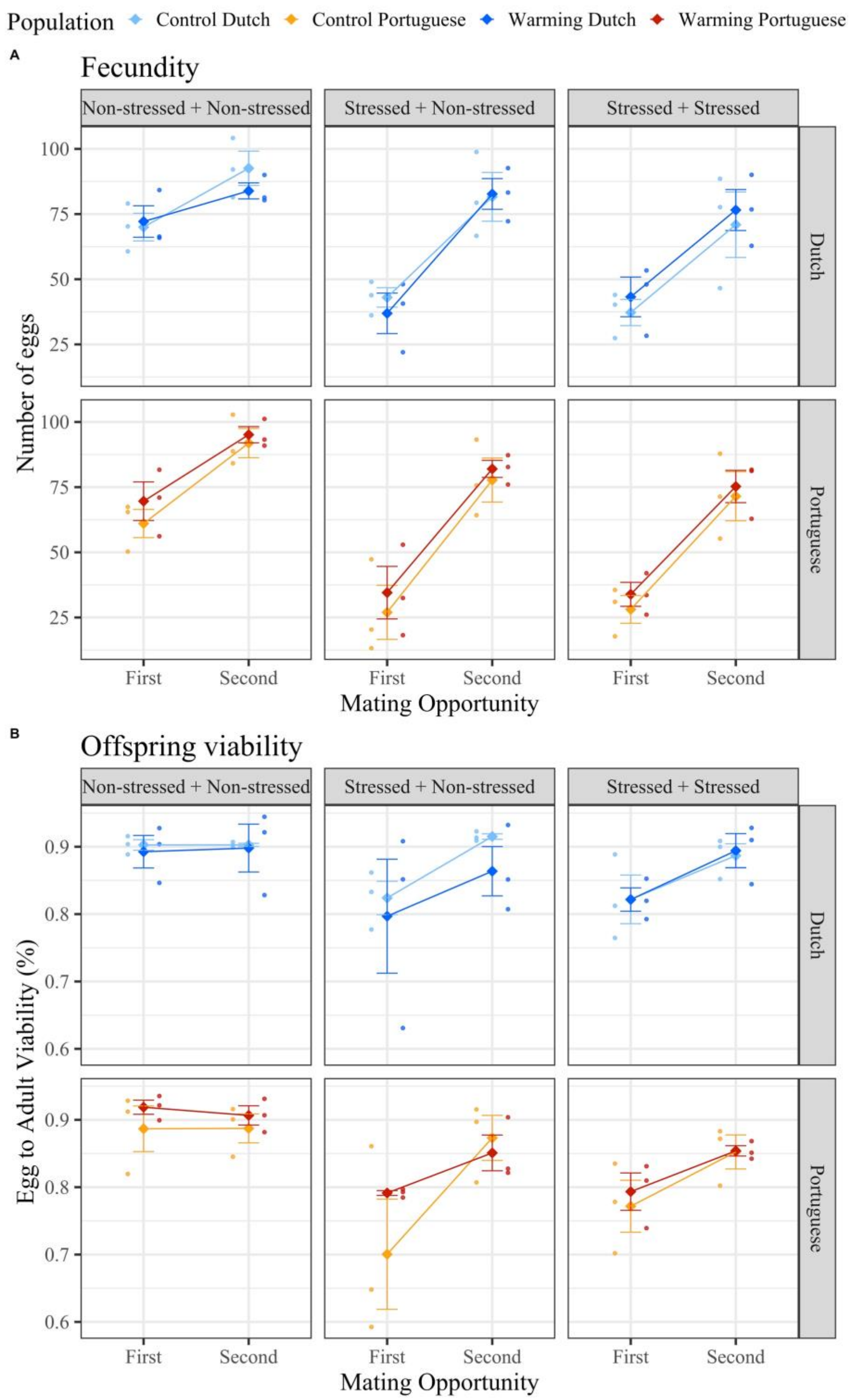


**Figure 2 – Effect of heatwave during male adulthood on female fertility.** A) Fecundity: number of laid eggs; B) Offspring viability: Ratio between the number of offspring and the number of laid eggs. Colder colours represent Dutch populations, while warmer ones represent Portuguese populations. Lighter tones represent the Control regime, while darker tones represent the Warming regime. The small circles represent the mean values of each replicate population, and the big diamonds represent the mean of the three replicate populations (±SE).
